# Detection of sex chromosomes in Tephritid pests using R-CQ and KAMY, two computational methods to enable species specific control

**DOI:** 10.1101/2023.10.27.564325

**Authors:** Dimitris Rallis, Konstantina T Tsoumani, Flavia Krsticevic, Philippos Aris Papathanos, Georgia Gouvi, Angela Meccariello, Kostas D Mathiopoulos, Alexie Papanicolaou

## Abstract

The detection and characterization of sex chromosome sequences is particularly important for major pest families, like the Tephritidae, whereas alternative pest management approaches, mainly involving male-only release programs, rely on the ability to target and manipulate sex-specific genomic regions, particularly those of the Y chromosome. However, resolving and detecting X and Y chromosome sequences at the chromosome level requires careful consideration of algorithmic outputs, especially in species where extensive sex chromosome markers are not available. Here, we present R-CQ and KAMY, two computational methods developed for the detection of sex chromosome-linked sequences. We evaluate their performance on newly generated chromosome-level assemblies of four important Tephritid pest species: *Ceratitis capitata, Bactrocera dorsalis, Bactrocera zonata* and *Anastrepha ludens*. By combining algorithmic predictions with a manual curation process, we assess the strengths and limitations of each method and provide a robust dataset of curated X- and Y-linked sequences. Overall, our results establish a framework for studying poorly characterized sex chromosome lineages and identifying sex-specific genomic regions, supporting the broader development of sex chromosome-based pest managements systems.

## Background

A common feature among many sexually dimorphic species is the presence of genetic sex determination signals, typically linked to specific chromosomes, known as sex-chromosomes (Bachtrog 2013; Bachtrog *et al*. 2014; Furman *et al*. 2020). In cases where sex chromosomes are observed as a pair, two systems exist, XY:XX where the Y chromosome is limited to males, and ZW:ZZ where females possess the sex-limited W chromosome (Bachtrog *et al*. 2014). In these systems, the Y (or W) chromosome often differs morphologically from its X (or Z) counterpart, making the pair heteromorphic. This heteromorphism is an outcome of suppressed recombination between the two sex chromosomes, a process that usually follows the acquisition of the sex determining function (Kent, Uzunović and Wright 2017; Furman *et al*. 2020; Guo *et al*. 2022). Over time, the sex-limited Y or W chromosome undergoes dramatic restructuring, characterized by the irreversible loss of gene content, the accumulation of repeats and the overall degeneration of the chromosome’s sequence (Bachtrog 2013; Mahajan and Bachtrog 2017).

Heteromorphic sex chromosomes, and specifically the Y, pose significant challenges for genomic projects due to their extensive repeat content and hemizygous regions, which complicate sequencing, assembly and scaffolding(Carey *et al*. 2022). It is common for the Y chromosome to be represented in small un-scaffolded contigs, mis-assembled or ever missing from the final genome releases (Hall *et al*. 2013). Recent advances in long-read sequencing and assembly technologies, coupled with chromatin-conformation capture methods like HiC, have provided high quality chromosome-level and haplotype-resolved assemblies that efficiently resolve repetitive and heterochromatic regions of the genome (Rhie *et al*. 2023; Li and Durbin 2024). Therefore, there is a growing potential in the application of these approaches for resolving the sequence and structure of repetitive and heterochromatic sex chromosomes.

The management of economically important pests, like many members of the Tephritidae family, may benefit from the careful reconstruction of their sex chromosome sequences and their subsequent genome editing with advanced CRISPR approaches (Meccariello *et al*. 2021; Nazarov *et al*. 2025). For sexually reproducing pests with heteromorphic sex chromosomes, as Tephritids, the sex-specific linkage of desired traits and the manipulation of sex chromosome inheritance are at the core of advanced management approaches (Meccariello *et al*. 2019, 2021). Sterile Insect Technique (SIT), is a long-established pest control strategy that relies on the selection, mass-rearing and sterilization of male-only individuals, which are then released into the field to suppress local populations by reducing their reproductive success (Dyck, Hendrichs and Robinson 2005). A critical requirement for SIT’s efficacy is the separation of males from females. Traditionally, this sex sorting has been achieved by introducing selectable genetic markers into the Y chromosome through chromosomal translocations. However, such methods are technically challenging, imprecise and largely limited to species with well-developed genetic tools. Recent advances in genome editing, particularly CRISPR/Cas technologies, offer a more targeted and efficient way of creating Genetic Sexing Strains (GSS), enabling the precise integration of selectable markers into non-recombining regions of the Y chromosome. Nevertheless, limitations on resolving, precisely detecting and annotating Y sequences may challenge the characterization of appropriate target sites. Therefore, the thorough reconstruction of sex chromosome sequences is crucial for considering candidate CRISPR target sites on the repetitive and heteromorphic Y, as the uniqueness and euchromatic regional context is considered pivotal for the efficient genome editing.

Crucially, in the absence of genetic markers for the identification and quality assessment of sex chromosome sequences in an assembly, as is the case in most non-model organisms, one must rely on computational methods. Several such methods have been described (Bernardo Carvalho and Clark 2013; Hall *et al*. 2013; Rangavittal *et al*. 2019) and successfully applied to Tephrtid sex chromosomes (Bayega *et al*. 2020). Nevertheless, their development and initial assessment was performed in model organisms with distinct genomic and sex chromosome characteristics. For example, *Drosophila melanogaster*, the primary model for Dipteran genomics, shares the same five large syntenic blocks known as the Muller elements, with Tephritids (Sved *et al*. 2016). However, different elements segregate as the sex chromosomes between these species, with the sex chromosome systems significantly varying in terms of content, origin and age (Vicoso and Bachtrog 2013; Meccariello *et al*. 2019).

To support the development of a generic approach for the application of SIT across different Tephritid pest species, long-read based male genome assemblies are now available for several economically important members of the family (Haig in preparation). In this study, we split the assemblies *of Ceratitis capitata, Bactrocera dorsalis, Bactrocera zonata and Anastrepha ludens* into their contig components to avoid cases of mis-scaffolding, and identify core regions that represent sex chromosome sequences.

We introduce and describe two computational pipelines, R-CQ and KAMY, designed for sex chromosome detection and curation. R-CQ employs a read alignment-based strategy, while KAMY leverages a kmer-based approach. We propose a manual curation methodology for assigning computational predictions to sex chromosomes, which is accompanied by cases of PCR validation. In this way we generate four robust core datasets of Tephritidae sex chromosome sequences. Finally, we present the utilities of KAMY in detecting Y-linked Tephritid genes, even in the absence of a high-quality Y chromosome reference sequence.

## Methods

### Species datasets & transcriptome assembly

The datasets used in this work include the male-derived chromosome level assemblies for *C. capitata* (GCA_905071925.1), *Bactrocera dorsalis* (GCA_037783525.1), *Bactrocera zonata* (GCA_037783105.1), *Anastrepha ludens* (GCA_037783455.1) and *Drosophila melanogaster* (GCA_000001215.4). For the application of sex chromosomes detection algorithms, the following short-read male and female-specific DNA sequencing datasets were used: *C. capitata* (ERR4026336, ERR4026337), *B. dorsalis* (SRR29323025, 2-BD_female_190927), *B. zonata* (SRR29319676, Bactrocera-Zonata_WGS_female_2-2659604_S1_L004_R1), *A. ludens* (SRR29323014, 4-AL_female_190916_L008) and *D. melanogaster* (ERR2163720, ERR2163721). For the transcriptome assembly of *A. ludens*, we used the Trinity assembler (Grabherr *et al*. 2011) with default parameters and used as input a concatenated read file containing the following testes-specific RNA-seq datasets: 31-51, 32-52, 33-53.

### Manual curation

During the manual curation process, the putatively X and Y contigs were reviewed from a single curator to assess the correctness of their assignment as sex chromosomes. This was done through observing the consistency and depth of male and female read-coverage across each contig on a genome browser (Diesh *et al*. 2023). We expect Y chromosome contigs to be consistently covered by male sequencing reads with ideally half the depth of the library, and partly from female sequencing reads due to the extensive presence of repeat on them that are also found in autosomal (or X) positions. Contigs were assigned as X chromosome when they were consistently supported by both female reads of the expected library depth and male reads with half the expected library depth. Genome browser tracks for both single- and multi-mappers were generated from short-read DNA sequencing using GSNAP(Wu Thomas D. and Reeder 2016). The results were further explored through graphs created in R (R Core Team 2022).

Examples of the different curation categories from *C. capitata* contigs are presented in Figure S. 14. Each putative sex chromosome-representing contig was inspected and assigned to one of the following categories:

- **Y**: assembly of Y regions covered predominantly by male reads with half the depth coverage of autosomal regions
- **X**: assembly of X regions consistently covered by male reads of half and female reads of normal autosomal depth
- **R:** putative repeat regions supported by extreme values of depth coverage for which the chromosomal origin cannot be confidently assigned
- **A**: autosomal region covered by male and female reads at the expected depth coverage

### Implementation of R-CQ algorithm

Reverse-CQ (R-CQ) algorithm is a re-implementation of the Chromosome Quotient method (Hall *et al*. 2013) that uses depth-coverage differences between male and female datasets to detect sex chromosome sequences. The uniquely mapping reads of each dataset need to be selected and transformed from a BAM into a lightweight BigWig file. Then, the algorithm will estimate the average library depth over a user-defined autosomal region that represents the diploid depth-coverage of each library, and use this to calculate a female normalization factor. Additionally, a repeat cutoff is calculated that excludes from the R-CQ calculation, regions with depth-coverage higher than twice the expected diploid. Sex chromosomes are detected in a genome by R-CQ through the calculation of the mean male-female ratio of depth-coverage values across non-overlapping sequence windows of predetermined size (100 bp by default). For cases where no female coverage exists for a window, the algorithm will set the female coverage of the window as 1 to avoid division by zero. The median ratio values are then calculated from all the windows of a sequence. Based on the principle of a male-only supported Y and a hemizygous X presence, we expect that Y contigs will have a large R-CQ value (ideally half the library depth), autosomes will have a ratio of 1 and X contigs will have an R-CQ of 0.5. The R-CQ values are converted to natural logarithms for visualization purposes through an R script and different chromosomal regions are initially assigned as follows; lnMedian(R-CQ) > 1 as putative Y sequences, 1 > lnMedian(R-CQ) > −0.4 as autosomal and lnMedian(R-CQ) < −0.4 as putative X-linked regions. In case additional contigs were observed near the X or Y threshold line, the respective settings were optimized for that species to include the whole putative X or Y cluster. The algorithm is freely available from https://zenodo.org/doi/10.5281/zenodo.10209595.

### Implementation of KAMY method

The “Kmer-counting Algorithm Meryl identifies the Y” KAMY method identifies Y-linked regions in an assembly by assessing their “breadth of coverage” (i.e. the proportion of a sequence) by sex-specific kmers, representing an alignment-free alternative. The method builds three distinct kmer databases (by default k=21) using Meryl (Rhie *et al*. 2020) from separate male and female genomic data: male, female and common. Kmers with less than two occurrences in the DNA-sequencing datasets are removed as sequencing errors. Subsequently, the breadth of coverage by either of the three kmer databases is assessed for each sequence of the query dataset and printed by the algorithm. Here, contigs were filtered as putative Y if they were covered by less than 90 % Common kmers, had at least 3 % higher Male coverage compared to Common and the (Male – Common) – (Female – Common) difference was higher than 5 %. Thresholds were determined empirically through the curation process and assessment of false positives. As KAMY checks the breadth of coverage by Male, Female and Common kmers across a sequence, it cannot be used to detect regions from the X, as such kmers are shared between males and females. For the transcriptome implementation, the direct output of Trinity was assessed. Sequences were initially size selected to be larger than 300 bp and filtered to contain transcripts with less than 20 % coverage by Common kmers. KAMY is freely available from https://zenodo.org/doi/10.5281/zenodo.10209589.

## Results

### Ceratitis capitata

The Mediterranean fruit fly *Ceratitis capitata* is a polyphagous species and one of the most destructive members of the family with extended presence across the globe (I. M. White and M. M. Elson-Harris 1992). The karyotype of Medfly consists of five pairs of autosomal chromosomes and a pair of heteromorphic sex chromosomes (Zacharopoulou *et al*. 2017). This species is characterized by elongated sex chromosomes, that appear as rod-like structures in the karyotype. Notably, X appears as the largest chromosome of the karyotype, while Y is characterized by two distinct chromosomal arms (Drosopoulou *et al*. 2017). The assembly of Medfly was based on PacBio CLR reads with 100× genome coverage and the detection of sex chromosomes was performed using a 26× Male and a 34× Female Illumina library. The employment of R-CQ and KAMY on the un-scaffolded Medfly genome led to the identification of 111 putative Y contigs, 96 commonly identified by both methods, 7 unique to R-CQ and 8 to KAMY (Figure 2, Table 1, Figure S. 1, Figure S. 2). The results were subjected to manual curation where contigs were marked as Y if they were supported predominantly of male reads with half the library depth, or Autosomal false positives if there was consistent coverage by female sequencing data. Importantly, the R-CQ method did not introduce any false positive autosomal contigs in the datasets in contrast to KAMY that contained 29 such regions in its predictions (Figure S. 2). To ensure the accuracy of manual curation, 10 Y curated contigs that contained computational gene prediction in the reference annotation were assessed through PCR, and all 10 were successfully amplified only from male DNA (Table 9). The total size of curated Y contigs summed up to ~27 Mb of Y sequence, while R-CQ and KAMY individually identified ~24 Mb and ~25 Mb of Y sequence respectively. The sum of false positive predictions was ~6.2 Mb identified only by KAMY, and ~4 kb of predictions comprised of Repeat regions (Figure 1, Table 1). Importantly, despite the high incidence of false positives introduced by KAMY, this method identified larger part of the Y compared to R-CQ, despite both methods having the same number of unique predictions (Table 1). Regarding the X chromosome, using R-CQ with a cutoff of lnMedian < −0.4 we detected 125 putative X contigs and 1 false positive autosome which accounts for ~52 Mb of curated X chromosome sequence (Figure 2, Figure S. 2, Table 7).

**Figure 1.**
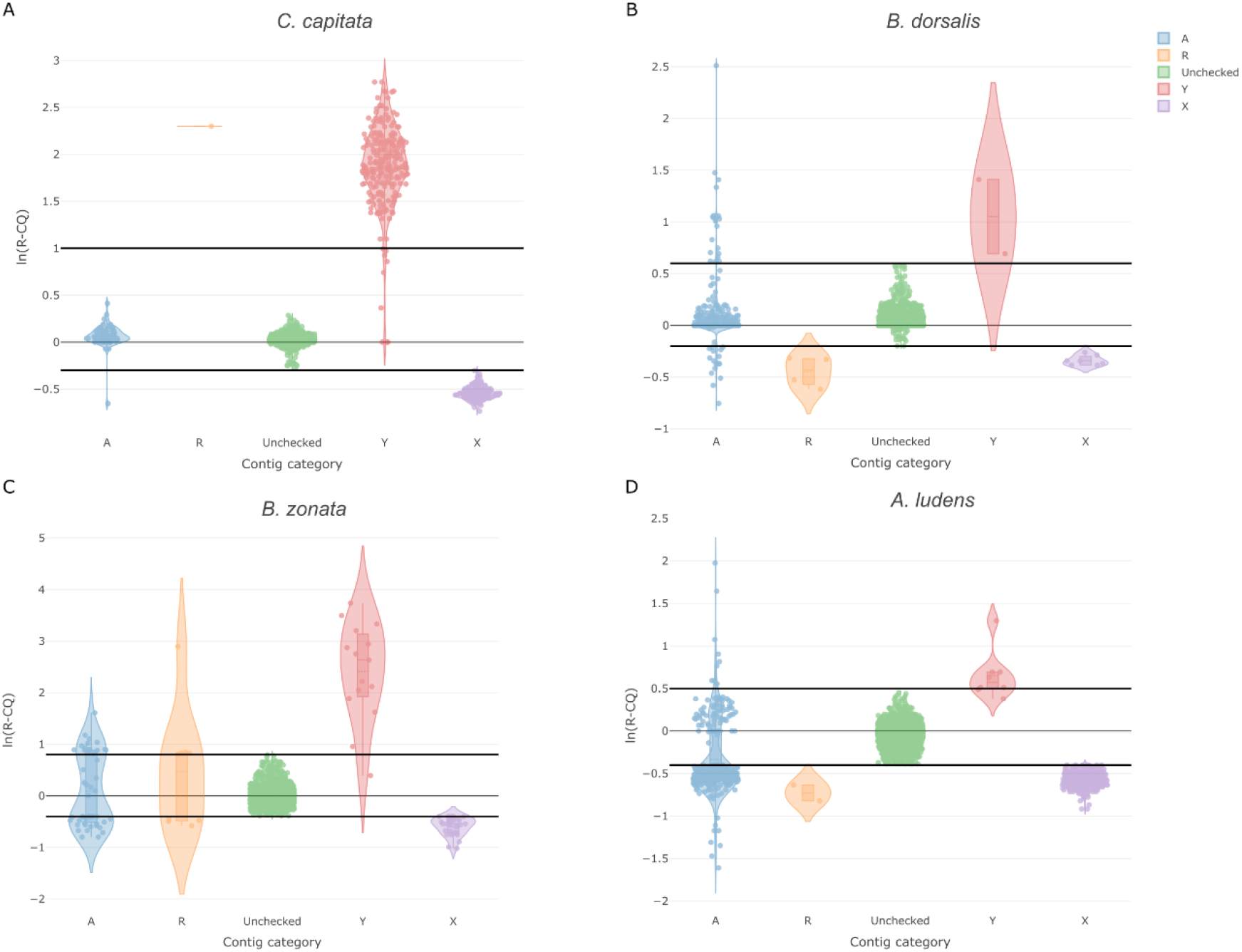
Curation of putative XY contigs from the four assessed Tephritids, relative to their ln(R-CQ) values.

**Figure 2.**
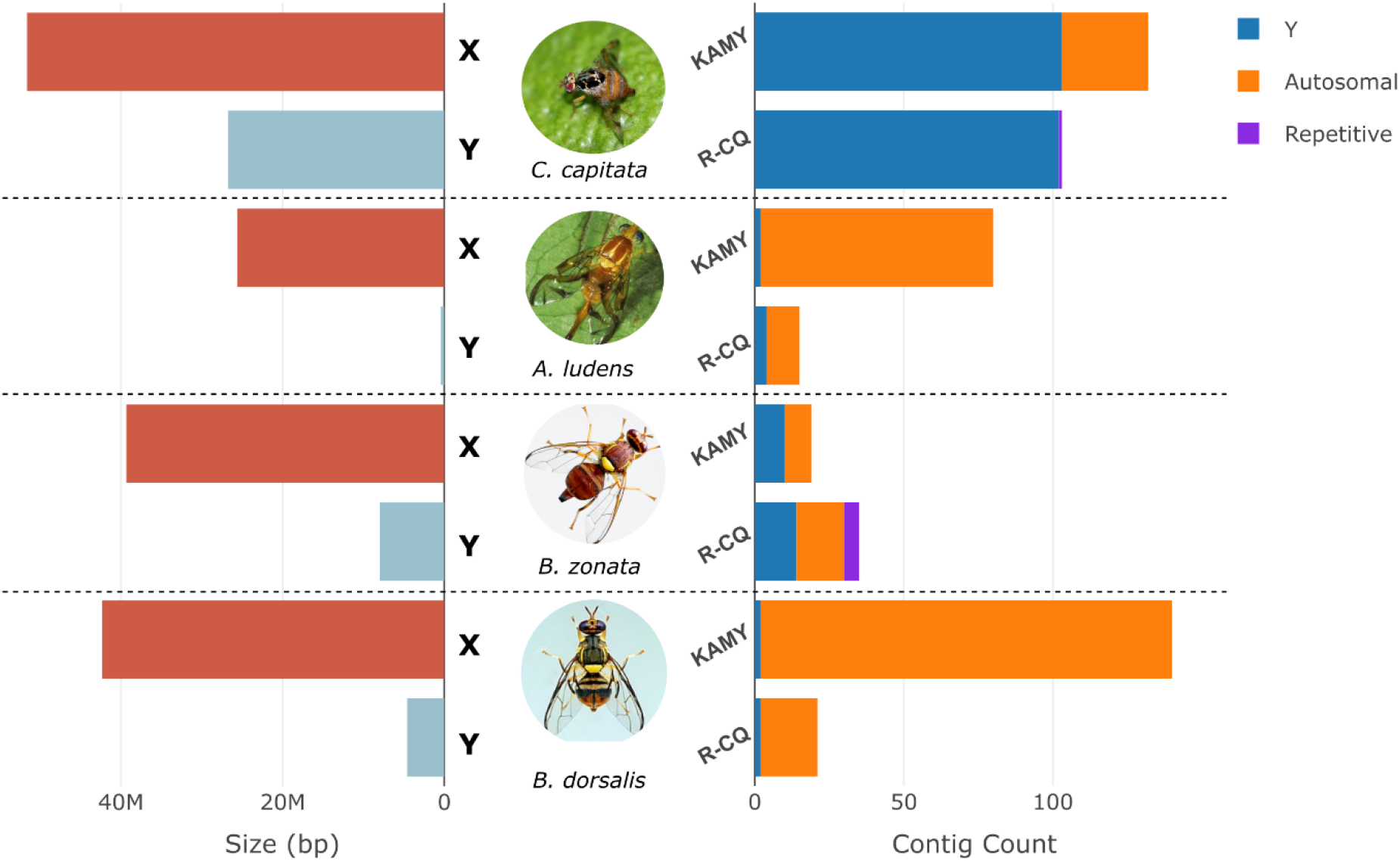
Collective results of sex chromosome identification against the long-read assemblies of Ceratitis capitata, Bactrocera dorsalis, Bactrocera zonata and Anastrepha ludens. The left panel represents the collective size of curated XY contigs of each species. The right panel shows the total Y predictions of R-CQ and KAMY and the respective curation.

Interestingly, the contig level assessment of the genome suggested that in the reference Medfly assembly, both X and Y contigs were scaffolded into a single hybrid scaffold. Specifically, 117 contigs identified as X-linked and 108 putative Y-linked contigs comprised the hybrid Scaffold_3. In addition, there were 3 putative Y and 8 putative X contigs misplaced in other scaffolds.

### Bactrocera dorsalis

The oriental fruit-fly *Bactrocera dorsalis* is considered one of the most notorious Tephritid pests due to its polyphagous and highly invasive character (Qin *et al*. 2018). The sex chromosomes of *B. dorsalis* comprise of a rod-like X chromosome and a dot-like Y, accompanied by five pairs of large autosomal chromosomes (Augustinos et al. 2014). The assembly of *B. dorsalis* was produced from PacBio CCS reads of 74x genome coverage and the detection of sex chromosomes was performed using a 16x Male and a 14x Female Illumina library. Splitting the chromosome-level scaffolds to their contigs revealed a highly contiguous assembly. Among them, R-CQ and KAMY predicted 21 and 169 contigs respectively, as putatively Y (Figure 2, Figure S. 3, Figure S. 4, Table 2). The manual curation suggested that only two of the predictions correspond to likely sequences from the Y chromosome. The remaining contigs, 19 R-CQ and 138 KAMY predictions were of autosomal origin, leading to an overall curated Y size of ~4.6 Mb, and ~56 Mb of false positives mainly contributed by KAMY (Figure S. 4, Table 2). The accuracy of manual curation for Y linkage was assessed through PCR, where 5 primer pairs amplifying regions of the curated Y contigs were designed, with all of them leading in the amplification of the desired product only from male DNA (Table 9). The high incidence of false positives by KAMY is considered a limitation related to the quality of the short-read DNA sequencing libraries and the representation of autosomal kmers in Male and Female databases. An autosomal contig is expected to be covered by the same percentage of Male and Female kmers, forming a diagonal line on the scatterplot. However, in cases where regions are stochastically underrepresented in the male compared to female sequencing data, this difference is reflected on the region being covered predominantly by female kmers, with such contigs found bellow the diagonal (higher proportion of coverage by female than male kmers). Similarly, the case of underrepresentation of autosomal regions in female sequencing data causes their coverage predominantly by male kmers, leading to their false categorization as Y-linked. As these discrepancies are randomly found across a contig, the calculation of depth-derived ratio of R-CQ is balanced by the consistent mapping of male and female data on the contig. For example, contig “scaffold_7.45” is 1.7 Mb in size and has 73.9 % coverage by male and 98.8 % by female kmers. However, the R-CQ value calculated from the Median of all the windows of the contig has a ratio of 1. As for the X chromosome, R-CQ predictions included 26 contigs, with the curation suggesting that 7 were putatively X-linked, with a total size of ~42 Mb, 15 false positive autosomal regions and 4 cases of unclassified Repeat contigs (Figure 2, Table 7, Table 8, Figure S. 4).

The scaffolds of *B. dorsalis* assembly were based on HiC proximity data. The contig-level detection of sex chromosomes indicated that the regions of the X and Y were again linked into a single hybrid XY scaffold. In contrast to Medfly, where many erroneous XY contig links were introduced, here the mis-scaffolding was caused by a single erroneous link between the edges of X and Y sequences. Surprisingly, no other fragments of the sex chromosomes were found in smaller regions of the assembly.

### Bactrocera zonata

The peach fruit-fly *Bactrocera zonata* is a major pest natively found in south Asia that attacks a variety of soft fruits (Zingore *et al*. 2020). The typical Tephritidae karyotype is also found in *B. zonata*, with five autosomal chromosomes and a pair of heteromorphic XY sex chromosomes (Yesmin and Hasanuzzaman 2021). In contrast to the sex chromosomes of *C. capitata* and *B. dorsalis, B. zonata* appears to contain a dot-like Y and a short X chromosome (Yesmin and Hasanuzzaman 2021). The assembly of *B. zonata* was generated using Nanopore High Accuracy model reads of 38x genome coverage and the detection of sex chromosomes was performed using a 45x Male and a 18x Female MGI DNBSEQ library. The contig analysis with R-CQ and KAMY suggested 29 and 15 predicted Y contigs respectively. Upon manual curation of the results, we concluded in 13 putative Y contigs; 10 were identified by R-CQ together with 14 autosomal false positives and 5 highly repetitive regions, while 9 were identified by KAMY with 6 autosomal false positives (Figure 2, Figure S. 5, Figure S. 6, Table 3). Among them, 6 contigs were commonly identified by both methods, R-CQ contributed 4 and KAMY 3 unique Y predictions (Table 3). The overall curated Y size reached ~7,9 Mb and since the different methods contributed unique Y contigs of relatively small size, the respective predicted Y size was ~7,6 Mb for R-CQ and ~7,3 Mb for KAMY (Figure S. 6, Table 3). False positives of R-CQ were also of small size, and despite the relatively large number, the total size of false predictions was ~1,1 Mb. On the other hand, KAMY had fewer false positives but of larger size, contributing ~1,4 Mb of false predictions. False positives were marked as Y due to the presence of regions covered predominantly by male-unique kmers, which however were not derived from the Y. In fact, the available *B. zonata* male libraries were sequenced at almost double the depth compared to female, which could possibly cause the underrepresentation autosomal regions and respective kmers in the female database. Overall, the sum of curated Y contig length was ~8 Mb with the respective false positive autosomal prediction summing to ~3.9 Mb. Regarding the X chromosome, R-CQ predicted 57 contigs as X, with manual curation suggesting that 28 were X-linked, 25 were false positives and 4 highly repetitive (Figure 2, Table 7, Table 8, Figure S. 6), with the curated size for X calculated at ~40 Mb (Figure 1).

The scaffolding of sex chromosomes in the *B. zonata* assembly resolved 5 contigs of the X as scaffold_8, 2 were linked in scaffold_1 and scaffold_5 and the rest comprised individual sequences. Two curated Y contigs were found to comprise scaffold_15, however the largest contiguous Y sequence was scaffold_9 that consisted of a single contig.

### Anastrepha ludens

The Mexican fruit-fly *Anastrepha ludens* is a polyphagous Tephritidae pest mainly found in Central and North America (Dupuis *et al*. 2019). The karyotype of *A. ludens* is characterized by the presence of five acrocentric autosomal chromosomes and a pair of heteromorphic XY sex chromosomes that include a highly degenerated Y (Zepeda-Cisneros *et al*. 2014). The assembly of *A. ludens* was generated using PacBio CLR reads of 30x genome coverage and the detection of sex chromosomes was performed using a 14x Male and a 11x Female Illumina library. Splitting the *A. ludens* scaffolds to their contig components indicated a highly fragmented assembly, over which we ran R-CQ and KAMY. R-CQ marked 15 contigs as putative Y with the respective number for KAMY summing to 80 putative Y contigs. The manual curation of the results suggested a total of 4 Y-linked contigs which summed to ~442 Kb of assembled Y sequence (Figure 1). Both methods commonly predicted 2 putative Y contigs and R-CQ uniquely contributed 2 additional Y contigs (Figure S. 8). The false positive predictions were found to be 11 contigs for R-CQ and 78 for KAMY, reaching a sum of ~6 Mb, mostly contributed by KAMY (Figure 2, Figure S. 8, Table 4, Figure S. 7, Figure S. 8). Regarding the X, 544 regions were marked as having higher depth coverage in females, with the curation suggesting that 342 were X-linked, 200 were autosomal false positive predictions and 2 contained highly repetitive sequences, the origin of which could not be confidently assigned (Figure 2, Table 7, Table 8). This resulted in a total of ~26 Mb of curated X sequence (Figure 1). The strengths and limitations of R-CQ and KAMY related to sequencing library depth are also observed here. Similar to the case of *B. dorsalis*, the low depth of sequencing results in many regions being underrepresented in either library, and the generation of falsely male- or female-specific kmers, which causes KAMY to include many autosomes in its prediction. On the other hand, R-CQ approach separates more efficiently Y contigs from false positive autosomes but struggles with clustering X from autosomal contigs. In the libraries used, the male diploid depth differs from haploid only by 7 reads, therefore the stochasticity of sequencing might randomly cause a lower male depth in regions, that deviates the Male/Female ratio toward the X-expected.

Despite the highly fragmented nature of the assembly, the HiC scaffolding managed to consecutively link 3 Y contigs in scaffold_14. No X sequences were found in scaffold_14 suggesting the absence of an XY mis-scaffolding in this assembly. Rather, contigs of the X were linked in many scaffolds, with the largest of them being scaffold_10.

### Testes transcriptome of A. ludens

The case of the highly fragmented *A. ludens* sex chromosome assembly is a common instance among non-model species and one of the major problems in the study of heteromorphic Y chromosome’s function in the context of its structure. An alternative approach that overcomes the need for long-read genomic data to provide Y transcribed elements is the generation of a de novo transcriptome assembly using RNAseq data. There, transcripts are assembled from RNA sequencing reads, independently of the quality of available genomic resources, which can provide insights into the transcribed elements of the Y. Since we expect for Y-linked genes to exert a male-specific role, a testes transcriptome assembly was produced here from tissue specific data of *A. ludens*. The candidate sequences were used as query in KAMY due to its ability to rapidly screen the candidate sequences and detect putative Y-derived transcripts from the already generated Male and Female kmer databases. A filtering threshold was set for candidate sequences to be covered by less than 20 % common kmers and to be larger than 400 bp. The results indicated four candidate Y transcripts, 7790_1, 7790_2, 90122 and 90852. Two of them, 7790_1 and 7790_2 were alternative isoforms of the same gene, with 7790_1 having 61 nucleotides more than 7790_2, while 90122 and 90852 contained different sequences. The two isoforms 7790_1 and 7790_2 had a length of 1,302 and 1,241 bp, while 90122 and 90852 were 1,167 bp and 1,165 bp respectively. We assessed the Y-linkage for every transcript of the four through PCR, and concluded to the presence of the genomic locus of 7790_1, 7790_2 and 90852 on the Y. For 90122 although the Male and Female PCR band slightly differed in size (Figure S. 14), we could not confidently assign the chromosomal origin therefore it was considered a false positive. Although the four selected transcripts had the highest coverage by male and lowest by female kmers, still not all of them could be amplified only from male DNA. This, in combination with the fact that many transcripts were covered by higher proportions of female than male kmers, suggest a library composition bias that is expected to induce false positives in the male-specific predictions (Figure S. 13).

### Drosophila melanogaster

The model species *Drosophila melanogaster* had been for years the reference point for exploring and understanding the genomes of Diptera species. Although the sex chromosomes of *D. melanogaster* are known to have a different origin that those of Tephritids (Vicoso and Bachtrog 2013), their presence and annotation in the reference assembly can serve as a positive control for validating the efficiency of R-CQ and KAMY since their sequence and elements are already studied and annotated. For that purpose, we used both the chromosome-levels scaffolds directly as well as the split contig version of *D. melanogaster*, applied KAMY and R-CQ and manually curated the results. Initially, both methods correctly marked the known Y scaffold as putative Y, with R-CQ containing 110 additional curated Y regions and 6 false positives. Similarly, KAMY included 132 curated Y regions and 5 false positives. Furthermore, the annotated-X scaffold was detected together with 41 smaller scaffolds, 7 small autosomal scaffolds and the scaffold that contained the mitochondrial sequence (Table 6, Table 7, Table 8, Figure S. 9, Figure S. 10). The scaffolded genome was also split to its contig component and methods were assessed for their ability to detect sex chromosome derived contigs. Out of the 207 contigs that comprised the Y scaffold, 125 were filtered out due to their size being smaller than 3 kb, 78 were predicted as Y while 4 contigs were not included in the predictions. Both methods commonly detected 63 of them, R-CQ uniquely contributed 3 contigs while KAMY had a higher sensitivity with 12 unique Y contig contributions (Table 5, Figure S. 11, Figure S. 12). The manual curation of missed Y contigs revealed that female data were consistently mapped in all of them, while in 3 the female coverage was double that of males, indicating a copy number variation in those regions between the two sexes and therefore a common occurrence of the sequence in both sex chromosomes. Assessing the sequence content of these contigs through BLAST revealed a composition by centromeric repeats, rDNA and transposon elements. Regions like these are known shared elements of X and Y chromosomes, explaining the common consistent presence of male and female data. On the side of X chromosome, out of the 10 contigs that comprised the annotated-X scaffold, 8 were effectively detected by R-CQ while the rest two were found to mainly contain incidents of highly repetitive regions that were excluded from the Male/Female ratio assessment due to R-CQ’s repeat cutoff (Figure S. 12, Table 7, Table 8).

## Discussion

In this study we present two computational methods, R-CQ and KAMY, for detecting sex chromosome sequences within genome assemblies. These approaches revisit well-established ratio-based and kmer-based strategies that have been used in sex chromosome-related studies for years (Bernardo Carvalho and Clark 2013; Hall *et al*. 2013). We applied these methods to four economically important Tephritid pest species; *Ceratitis capitata, Bactrocera dorsalis, Bactrocera zonata* and *Anastrepha ludens*, by splitting their chromosome-level assemblies into their contig components and manually curating the extent and scaffolding quality of sex chromosome sequences. Additionally, we used the chromosome-level assembly of *D. melanogaster* as a control to validate the competence of our methods. R-CQ and KAMY were built towards a flexibility in usage and broad applicability, ensuring their suitability for diverse datasets and different queries. R-CQ uses lightweight binary BigWig files as input form the mapping of short-reads to the query, screens the sequence for Male/Female depth ratios in a window-based manner, while performing library depth normalization and repeat exclusion, providing the detection of sequencing depth variability between male and female data that might be linked to X and Y unique regions. KAMY generates a set of kmer databases from male and female DNA sequencing reads, screens the query for breadth of coverage by either Male, Female or Common kmers, allowing the detection of regions covered predominantly by male kmers which might reflect Y sequences.

The combined application of both R-CQ and KAMY over the assessed assemblies resulted in the overall detection of curated Y (*C. capitata*; ~47 Mb, *B. dorsalis*; ~4.6 Mb, *B. zonata*; ~7.9 Mb, *A. ludens*; ~442 kb) and X sequences (*C. capitata*; ~52 Mb, *B. dorsalis*; ~42 Mb, *B. zonata*; ~39 Mb, *A. ludens*; ~26 Mb) from all four species (Figure 1, Figure 2). By using the same filtering parameters against all datasets, KAMY was found to underperform in many cases, introducing many false positive autosomal contigs in the results. Despite this limitation, both methods contributed unique Y predictions in each dataset, with the exception of *the B. dorsalis* Y (Figure S. 2, Figure S. 4, Figure S. 6, Figure S. 8). In addition, the implementation of KAMY against the transcriptome dataset of *A. ludens* demonstrated an effective and low-cost alternative for identifying Y-linked transcribed elements. In fact, the newly identified *A. ludens* Y-linked genes were the ones marked by KAMY as having the highest coverage by male and lowest coverage by female kmers, suggesting Y-linkage. Additional Y-linked genes might exist in the dataset that were filtered out by the coverage thresholds, as many transcripts existed with higher male kmer coverage (Figure S. 13). However, the fact that many transcripts of the dataset were covered predominantly by female kmers (Figure S. 13) suggests an uneven representation of genomic regions in the sequencing data. With higher quality sequencing libraries, we expect a reduction in false positives and an improved ability of KAMY to accurately detect additional Y-linked transcripts. Nevertheless, our results suggest KAMY’s flexibility across different query datasets and its capacity to rapidly identify Y-linked sequences.

In all cases, we observed that the detection methods highly relied on the depth and quality of DNA sequencing libraries. Although R-CQ includes a female depth normalization step to correct for differences between male and female libraries, sequencing depth was found to be a limiting factor in the specificity of the method, as many autosomal false positives were introduced, especially over the detection of X-linked sequences. Based on R-CQ, the mean depth in the normalizer autosomal contig is 26× and 34× for Medfly male and female libraries respectively, and the Male/Female ratio calculation from these libraries results in three distinct clusters for X, Y and autosomes. Conversely for *A. ludens*, the mean depth values were found to be 14× for males and 11× for females, which given the stochasticity of sequencing, might cause the underrepresentation of genomic regions in the sequencing dataset, leading to discrepancies in the Male/Female ratio calculation. As a result, many autosomal contigs might deviate from the expected ln(R-CQ) ratio of zero, being marked as candidate X or Y sequences from the analysis (Figure 2). Nevertheless, even in this case the majority of autosomal contigs were clustered in an expected ln(R-CQ) of zero, suggesting that even under these circumstances the depth normalization step and the window-based R-CQ calculation can calibrate variations between the two different libraries.

Since both KAMY and R-CQ rely on distinct computational approaches for detecting sex chromosomes, variability in the performance is expected depending on the quality and composition of the input genomic data. Therefore, KAMY and R-CQ should be considered complementary approaches rather than comprehensive solutions. KAMY struggled with the underrepresentation of sequences in the Male and Female databases of *B. dorsalis* while low sequencing depth in *A. ludens* led to R-CQ underperformance in identifying X-linked contigs. Manual curation is therefore important to assess how male and female sequencing reads support each candidate XY region, and understand putative problems with the genomic datasets that might introduce false positive results. Although time-consuming, this step helps refine the dataset by removing false positives, thereby increasing the precision and robustness of sex chromosome detection and revealing the interaction between dataset quality and method performance. The fact that computational methods implement models to interpret a biological state should always be acknowledged and approached critically, especially when it comes to heterochromatic elements with single-copy state like sex chromosomes, where sequencing methods are known to underperform.

Throughout our analysis, we also emphasize the importance of assessing sex chromosomes in the contig level. Chromatin conformation capture technologies, like HiC, provide proximity information for genomic regions, aiding to the linkage of assembly regions to chromosome-like structures (Haig in preparation). However, we observed that discrepancies might occur in the case of sex chromosomes, the nuclear interactions of which are mostly unexplored. Medfly is a characteristic example, as the XY chromosomes of that species were mis-scaffolded in a single scaffold, with the cause of that being still unclear. However, karyotypic studies of Medfly in trichogen cells describe the sex chromosomes as a heterochromatic network (Bedo 1987) and detect on them clusters of ribosomal genes (Bedo and Webb 1989). These characteristics suggest spatial interactions between X and Y chromosomes, which may complicate the resolution of sex chromosome proximal regions, causing the false linkage of X and Y contigs. A similar situation was observed in *B. dorsalis*, where X and Y chromosomes were again linked in a single scaffold, yet the mis-scaffolding had occurred to a smaller extent and was attributed to a single link between the terminal regions of the X and Y chromosomes.

Despite challenges with the scaffolding of sex chromosomes, long-read sequencing and assembly technologies managed to resolve to different extent the sequences of the highly repetitive and heterochromatic Tephritidae Y chromosomes. Different long-read sequencing platforms were employed for producing the genomes of the assessed species. The assemblies of *C. capitata* and *A. ludens* were based on the error-prone PacBio CLR reads with an error rate of 11-15 % (Korlach), the genome of *B. zonata* was generated based on high-accuracy model of Nanopore reads with > 99 % accuracy (Kim et al. 2024), while that of *B. dorsalis* relied on HiFi CCS PacBio reads with a > 99 % accuracy (Korlach). As expected, the difference between platforms is reflected in the quality of the Y chromosome assemblies. Although *A. ludens* and *C. capitata* genomes were both assembled using PacBio long-reads, they differ significantly in sequencing depth, with the former being assembled with 30x and the latter with 100x CCS coverage. It is therefore not surprising that the assembler struggled to resolve the hemizygous Y chromosome of *A. ludens* from the error-prone CCS reads, while the significantly higher depth of *C. capitata* was adequate to produce well-assembled Y sequences. The advances of the PacBio output are obvious in the case of *B. dorsalis*, where the HiFi CCS reads from a 74x library depth produced highly contiguous sequences of the Y chromosome. On the other hand, Nanopore sequencing performed equally well in resolving the *B. zonata* Y chromosome under a significantly lower sequencing depth of 38x. However, a small underperformance was observed among the putative Y-linked contigs, as the predictions included regions of extremely high coverage, likely representing common autosome-Y repetitive elements. These contigs were initially classified as putative Y-linked based on the Male/Female coverage but could not be confidently assigned as Y-linked during the curation process due to the consistent support by female data. However, we can assume from the highest occurrence of male reads in these regions that these might represent a cluster of common repeats shared with the Y chromosome, that were not resolved appropriately due to the higher error inherent to Nanopore sequencing. Similar patterns were also found in the *C. capitata* assembly that was also based on error-prone reads, albeit to a smaller extent.

The independent evolution of sex chromosome systems across diverse linages has been a fascinating field for genome biologists. These systems, particularly the XY (or ZW) configuration, not only offer insights into chromosome evolution but also present unique opportunities for applied biology. The Y chromosome, for example, is found in a single copy only in males and often harbors male-specific sequences. This makes it a powerful, sex-specific genomic platform for the development of pest management strategies that manipulate reproductive dynamics. One such strategy is the long-standing Sterile Insect Technique (SIT) that involves releasing sterilized male insects into the wild to reduce pest populations. A critical component of SIT is the use of Genetic Sexing Strains (GSS), in which selectable markers are linked to the Y chromosome to produce male-only cohorts for release. For years, GSSs were created through chromosomal translocations, a technically challenging and imprecise approach. Recent advances in molecular tools, particularly CRISPR/Cas systems, offer a more targeted and efficient alternative for developing GSS. Beyond simple marker integration, the use of CRISPR has also been proposed to bias sex chromosome transmission by shredding the X chromosome in the male germline, leading to male-biased progeny and reduced reproduction potential in wild populations (Meccariello *et al*. 2021). Importantly, the content of the Y chromosome, not just its presence, is crucial to SIT applications. Y-linked genes that contribute to male fertility or mating competitiveness can directly impact the effectiveness of released males. Therefore, a deeper understanding of the Y chromosome structure and gene content is essential for the optimization of SIT-based pest control. Complementary to this, advanced synthetic gene drive systems like precision guided SIT (pgSIT) can target male-limited fertility genes for precise CRISPR-mediated genetic sterilization of released insects (Kandul *et al*. 2019)(Sima in preparation). Moreover, the discovery of the Tephritid Maleness factor has opened up avenues for conditional sex conversion systems that further expand the genetic control toolkit (Meccariello et al. 2019).

The emergence of third-generation sequencing technologies now allows for the high-resolution assembly of repetitive and heterochromatic regions, including sex chromosomes. Leveraging these advances, our study explores the recently released genomes of four Tephritidae pest species, *Ceratitis capitata, Bactrocera dorsalis, Bactrocera zonata* and *Anastrepha ludens*, and introduces two computational pipelines, R-CQ and KAMY, designed to detect and curate sex chromosome sequences. Together, our methods provide a framework for detecting and evaluating X- and Y-linked elements. We expect this work to support the broader implementation of GSS in SIT programs and to enable comparative studies on sex chromosome structure, function and evolution.

## Supporting information

Supplementary matterial

## Acknowledgements

Funding for this research was provided by the European Union’s Horizon Europe Research and Innovation Programme REACT (Grant agreement 101059523). The publication of this work is supported by the International Atomic Energy Agency’s Coordinated Research Project “Generic approach for the development of genetic sexing strains for SIT applications”.

## Disclosure

The authors declare no competing interests.

---

*Table 1 Curation of R-CQ and KAMY output over C. capitata split genome*.

*Table 2 Curation of R-CQ and KAMY output over B. dorsalis split genome*.

*Table 3 Curation of R-CQ and KAMY output over B. zonata split genome*.

*Table 4 Curation of R-CQ and KAMY output over A. ludens split genome*.

*Table 5 Curation of R-CQ and KAMY output over D. melanogaster split genome*.

*Table 6 Curation of R-CQ and KAMY output over D. melanogaster scaffolded genome*.

*Table 7 Curations and total size of R-CQ X chromosome outputs against the genomes of assessed species. Abbreviations stand for Ceratitis capitata (Cc), Bactrocera dorsalis (Bd), Bactrocera zonata (Bz), Anastrepha ludens (Al) and Drosophila melanogaster (Dm)*.

*Table 8 Primers used for PCR validation of Y-linkage from C. capitata contigs.*

*Table 8 Primers used for PCR validation of Y-linkage from B. dorsalis contigs*.

*Figure S. 1 Scatterplot of male VS female kmer coverage of C. capitata contigs as identified by KAMY. A) kmer coverage of total C. capitata contigs. B) Curation-based classification of total C. capitata contigs in relation to kmer coverage values. C) kmer coverage of putative Y C. capitata contigs. D) Curation of putative Y contigs identified by KAMY*.

*Figure S. 2 Violin plots of ln(R-CQ) values for C. capitata contigs, horizontal line indicates the ln=1 and ln= −0.3 cutoff used for indicating Y and X contigs respectively. A) Contigs are separated into four size-based quartiles. B) Distribution of ln(R-CQ) values over different curation categories*.

*Figure S. 3 Scatterplot of male VS female kmer coverage of B. dorsalis contigs as identified by KAMY. A) kmer coverage of total B. dorsalis contigs. B) Curation-based classification of total B. dorsalis contigs in relation to kmer coverage values. C) kmer coverage of putative Y B. dorsalis contigs. D) Curation of putative Y contigs identified by KAMY*.

*Figure S. 4 Violin plots of ln(R-CQ) values for B. dorsalis contigs, horizontal line indicates the ln=0.6 and ln= −0.2 cutoff used for indicating Y and X contigs respectively. A) Contigs are separated into four size-based quartiles. B) Distribution of ln(R-CQ) values over different curation categories*.

*Figure S. 5 Scatterplot of male VS female kmer coverage of B. zonata contigs as identified by KAMY. A) kmer coverage of total B. zonata contigs. B) Curation-based classification of total B. zonata contigs in relation to kmer coverage values. C) kmer coverage of putative Y B. zonata contigs. D) Curation of putative Y contigs identified by KAMY*.

*Figure S. 6 Violin plots of ln(R-CQ) values for B. zonata contigs, horizontal line indicates the ln=0.8 and ln= −0.4 cutoff used for indicating Y and X contigs respectively. A) Contigs are separated into four size-based quartiles. B) Distribution of ln(R-CQ) values over different curation categories*.

*Figure S. 7 Scatterplot of male VS female kmer coverage of A. ludens contigs as identified by KAMY. A) kmer coverage of total A. ludens contigs. B) Curation-based classification of total A. ludens contigs in relation to kmer coverage values. C) kmer coverage of putative Y A. ludens contigs. D) Curation of putative Y contigs identified by KAMY*.

*Figure S. 8 Violin plots of ln(R-CQ) values for A. ludens contigs, horizontal line indicates the ln=0.5 and ln= −0.4 cutoff used for indicating Y and X contigs respectively. A) Contigs are separated into four size-based quartiles. B) Distribution of ln(R-CQ) values over different curation categories*.

*Figure S. 9 Scatterplot of male VS female kmer coverage of D. melanogaster scaffolds as identified by KAMY. A) kmer coverage of total D. melanogaster scaffolds. B) Curation-based classification of total D. melanogaster scaffolds in relation to kmer coverage values. C) kmer coverage of putative Y D. melanogaster scaffolds. D) Curation of putative Y contigs identified by KAMY*.

*Figure S. 10 Violin plots of ln(R-CQ) values for D. melanogaster scaffolds, horizontal line indicates the ln=0.8 and ln= −0.2 cutoff used for indicating Y and X scaffolds respectively. A) Scaffolds are separated into four size-based quartiles. B) Distribution of ln(R-CQ) values over different curation categories*.

*Figure S. 11 Scatterplot of male VS female kmer coverage of D. melanogaster contigs as identified by KAMY. A) kmer coverage of total*

*D. melanogaster contigs. B) Curation-based classification of total D. melanogaster contigs in relation to kmer coverage values. C) kmer coverage of putative Y D. melanogaster contigs. D) Curation of putative Y contigs identified by KAMY*.

*Figure S. 12 Violin plots of ln(R-CQ) values for D. melanogaster contigs, horizontal line indicates the ln=0.6 and ln= −0.3 cutoff used for indicating Y and X contigs respectively. A) Contigs are separated into four size-based quartiles. B) Distribution of ln(R-CQ) values over different curation categories*.

*Figure S. 13 Scatterplot of male VS female kmer coverage of A. ludens testes transcripts as identified by KAMY. A) kmer coverage of total A. ludens testes transcripts. B) kmer coverage of putative Y A. ludens testes transcripts*

*Figure S. 14 PCR validation of the putative Y transcripts of A. ludens 7790, 7790_2, 90122 and 90852. Line M and F contain the male and female specific amplification product while red M contains the molecular weight Marker*.

*Figure S. 15 Screen snips of the male and female coverage plots from the Genome Browser of C. capitata. A) Example of a Y contig. B) Example of a false positive Autosomal contig. C) Example of a contig of the R (repeats) category. D) Example of an X contig*.

## Notes

https://zenodo.org/doi/10.5281/zenodo.10209595

https://zenodo.org/doi/10.5281/zenodo.10209589

### Competing Interest Statement

The authors have declared no competing interest.

### Summary of Updates

The long-read based assemblies of four Tephritidae pest species were assessed with the KAMY and R-CQ sex chromosome detection methods, in addition to the Diptera model Drosophila melanogaster as a demonstration of the interaction between genomic data and sex chromosome detection methods in long-read based assemblies. The results were subjected to manual curation and the identified regions are reported here. In addition, detection and curation of X chromosome sequences was performed.

