## Supplementary matterial for "Detection of sex chromosomes in Tephritid pests using R-CQ and KAMY, two computational methods to enable species specific control"

### Supplementary figures

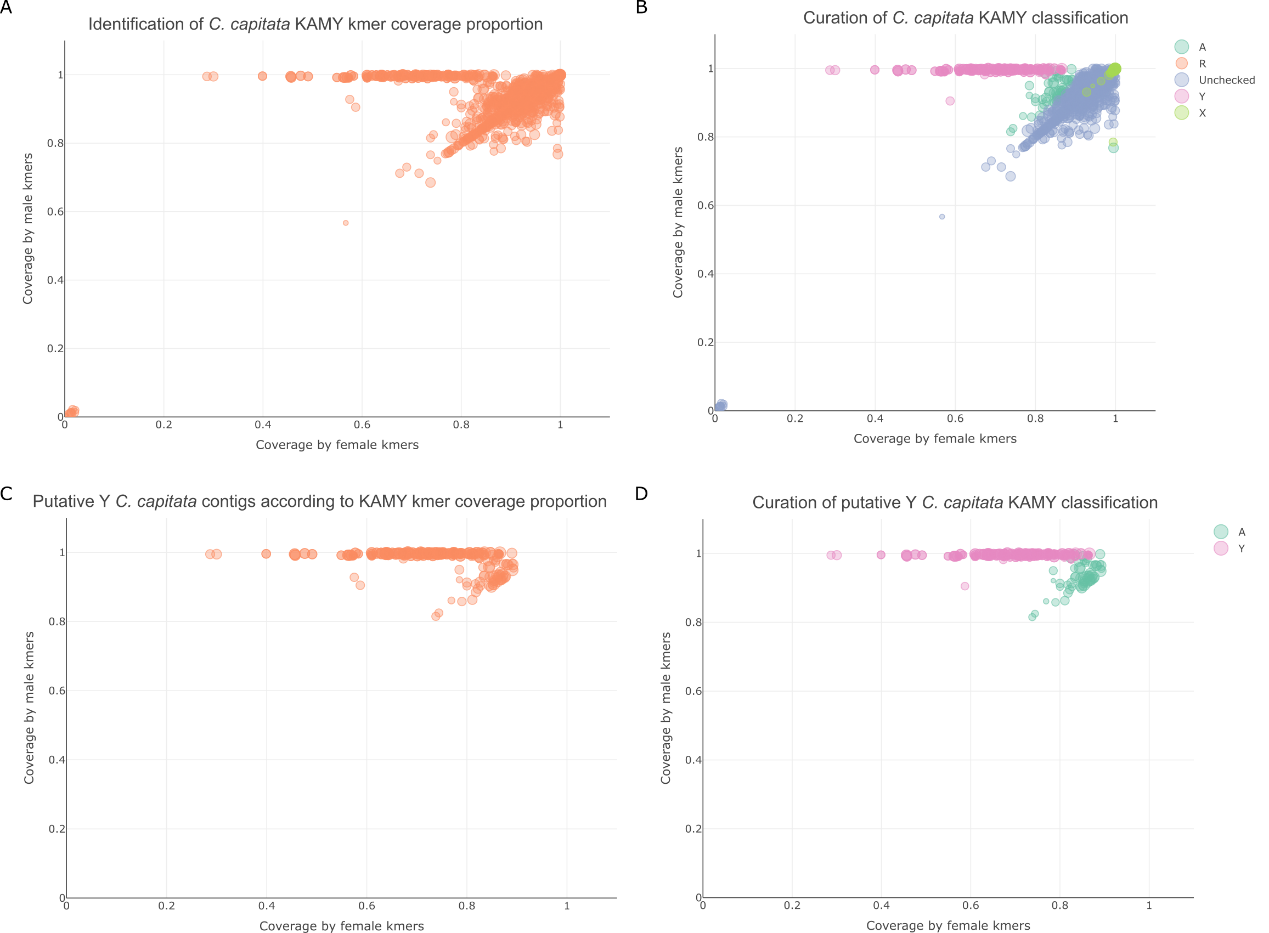

Figure S. 1 Scatterplot of male VS female kmer coverage of C. capitata contigs as identified by KAMY. A) kmer coverage of total C. capitata contigs. B) Curation-based classification of total C. capitata contigs in relation to kmer coverage values. C) kmer coverage of putative Y C. capitata contigs. D) Curation of putative Y contigs identified by KAMY.

Figure S. 2 A) Violin plot of ln(R-CQ) values for C. capitata, with contigs separated into four size-based quartiles. Horizontal lines indicate the ln=1 and ln= -0.3 cutoff used for indicating Y and X contigs respectively. B) Size distribution of curation-based Y and false positive contigs across R-CQ and KAMY predictions for C. capitata.

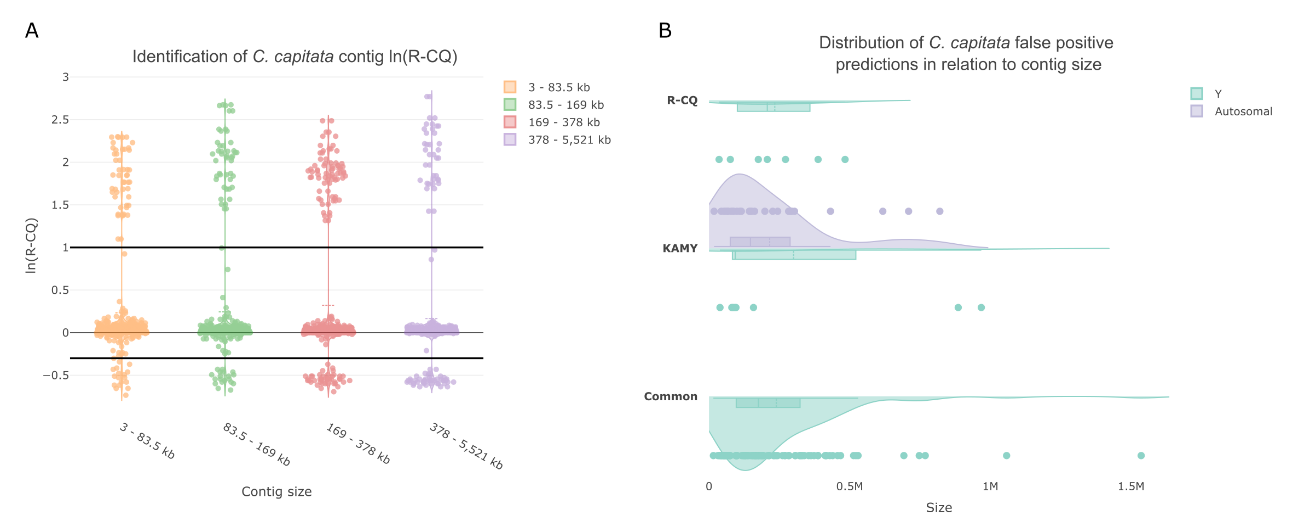

Figure S. 3 A) Violin plot of ln(R-CQ) values for B. dorsalis, with contigs separated into four size-based quartiles. Horizontal lines indicate the ln=0.6 and ln= -0.2 cutoff used for indicating Y and X contigs respectively. B) Size distribution of curation-based Y and false positive contigs across R-CQ and KAMY predictions for B. dorsalis.

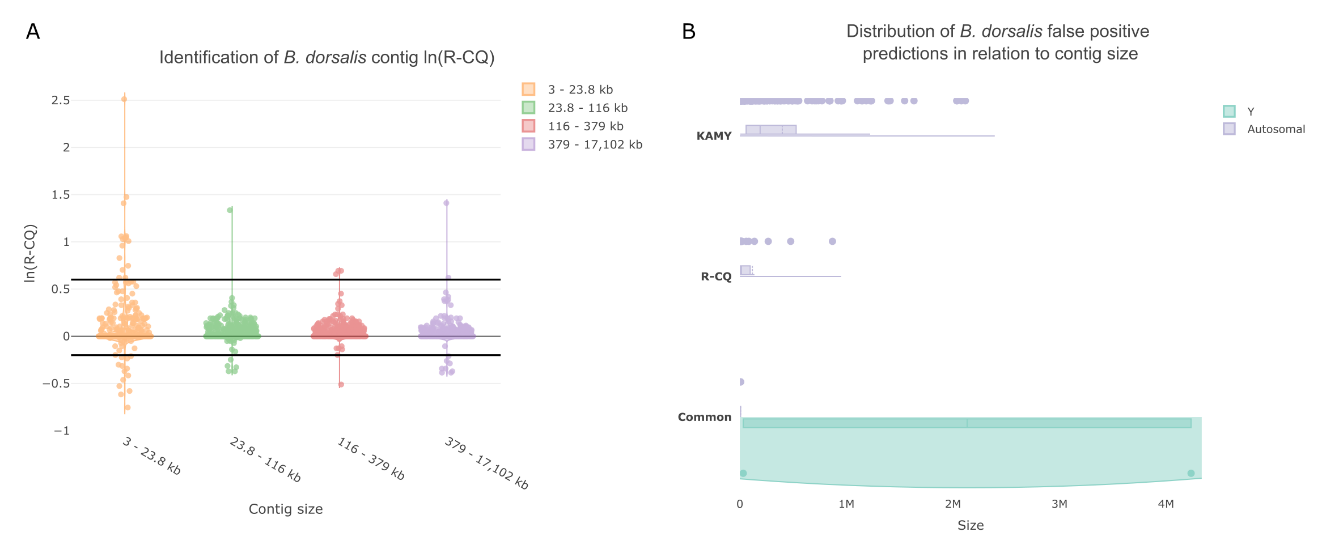

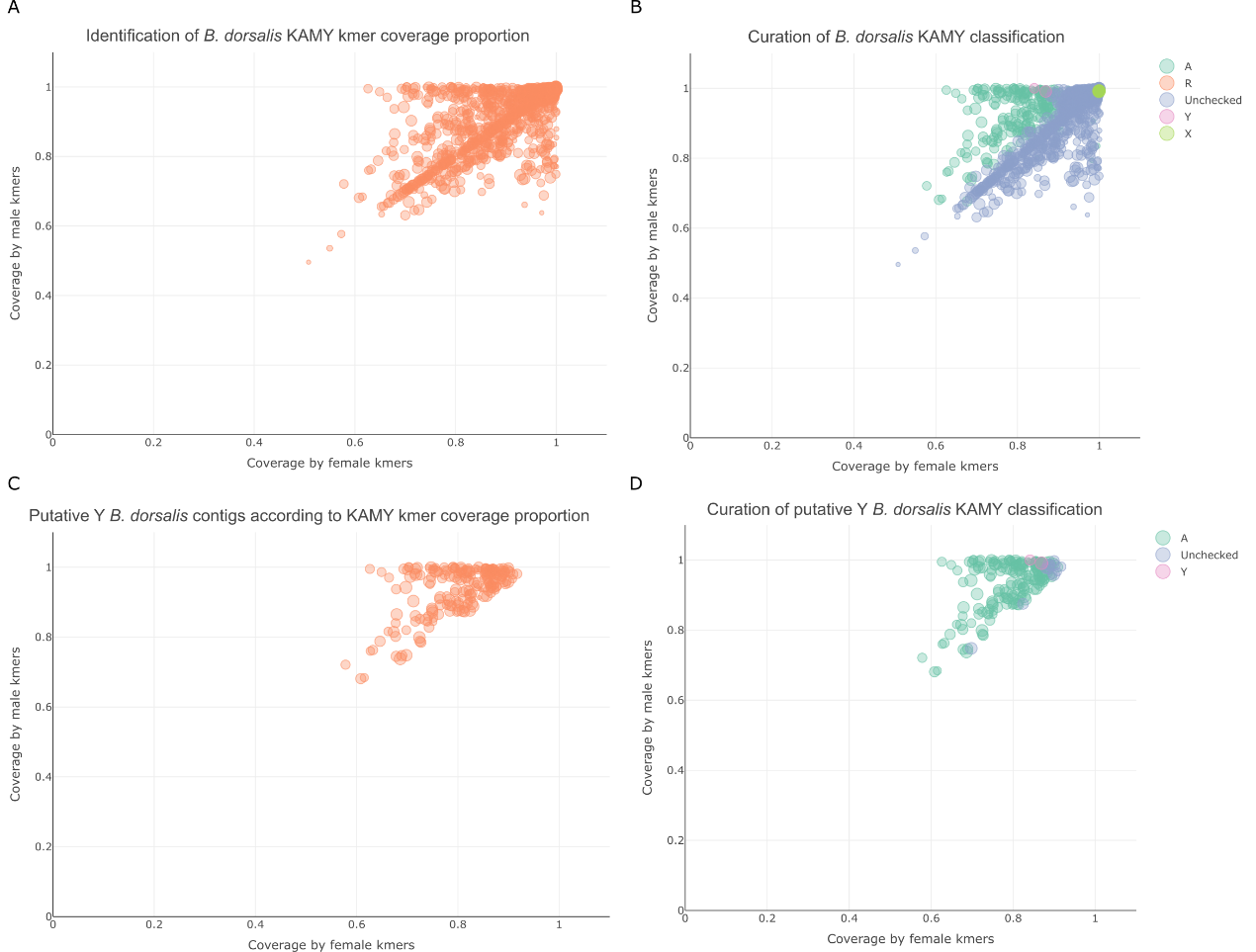

Figure S. 4 Scatterplot of male VS female kmer coverage of B. dorsalis contigs as identified by KAMY. A) kmer coverage of total B. dorsalis contigs. B) Curation-based classification of total B. dorsalis contigs in relation to kmer coverage values. C) kmer coverage of putative Y B. dorsalis contigs. D) Curation of putative Y contigs identified by KAMY.

Figure S. 5 A) Violin plot of ln(R-CQ) values for B. zonata, with contigs separated into four size-based quartiles. Horizontal lines indicate the ln=0.8 and ln= -0.4 cutoff used for indicating Y and X contigs respectively. B) Size distribution of curation-based Y and false positive contigs across R-CQ and KAMY predictions for B. zonata.

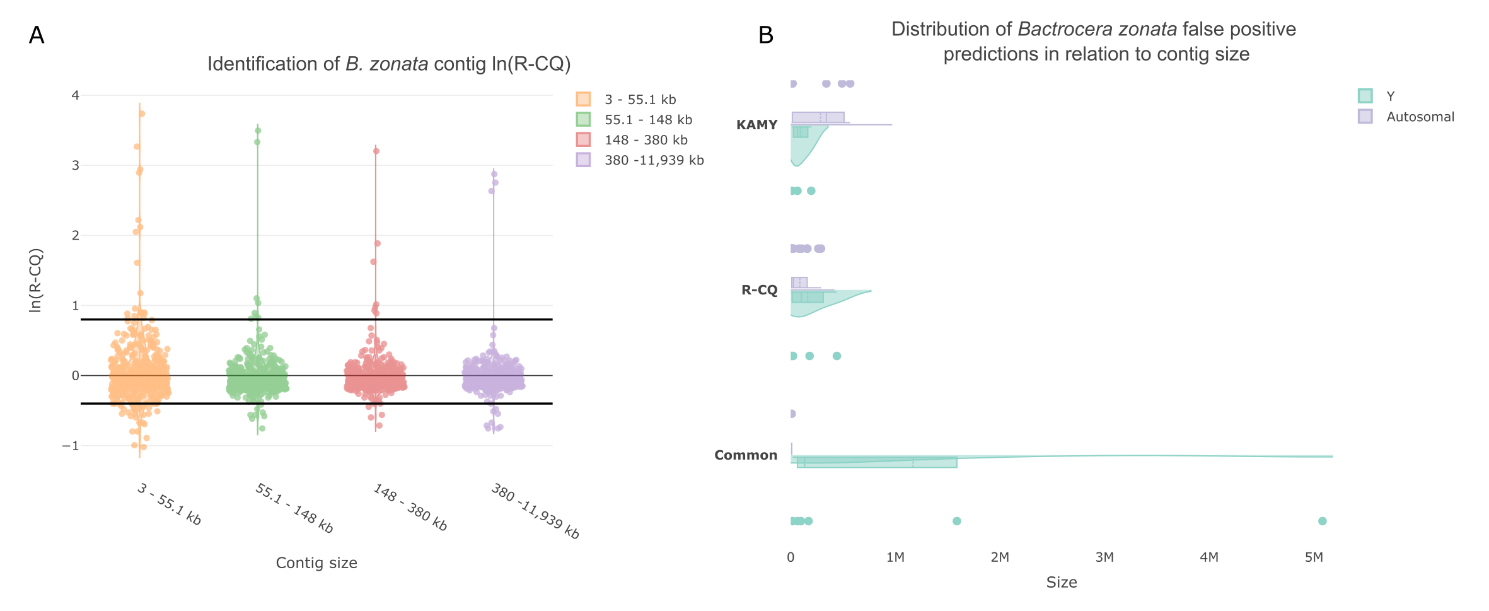

Figure S. 6 Scatterplot of male VS female kmer coverage of B. zonata contigs as identified by KAMY. A) kmer coverage of total B. zonata contigs. B) Curation-based classification of total B. zonata contigs in relation to kmer coverage values. C) kmer coverage of putative Y B. zonata contigs. D) Curation of putative Y contigs identified by KAMY.

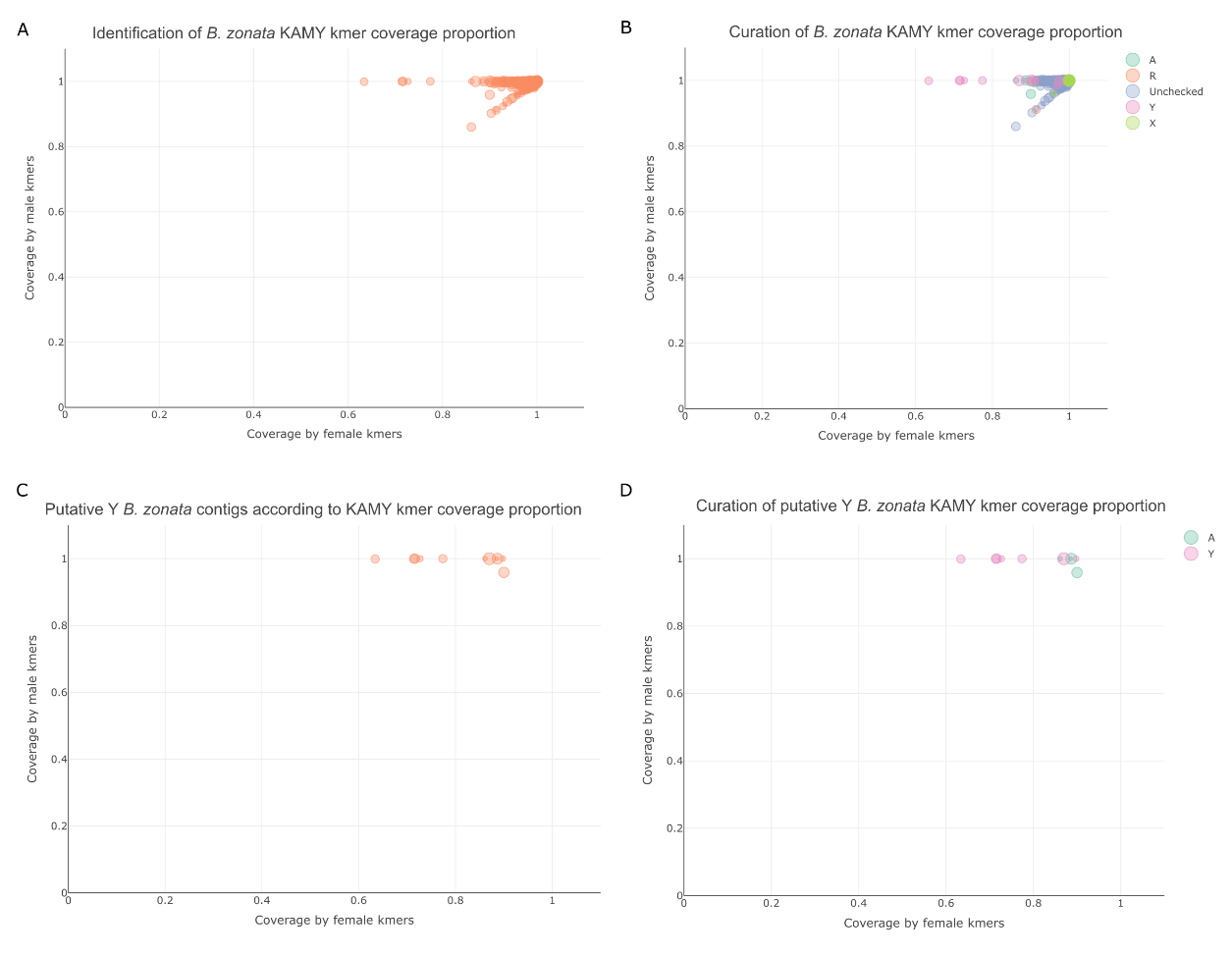

Figure S. 7 A) Violin plot of ln(R-CQ) values for A. ludens, with contigs separated into four size-based quartiles. Horizontal lines indicate the ln=0.5 and ln= -0.4 cutoff used for indicating Y and X contigs respectively. B) Size distribution of curation-based Y and false positive contigs across R-CQ and KAMY predictions for A. ludens.

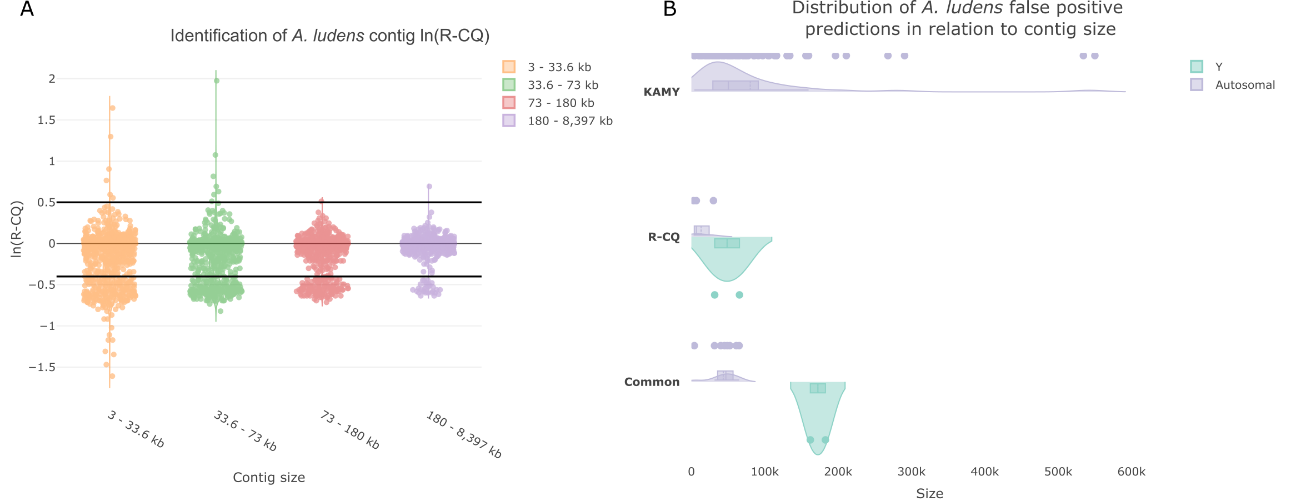

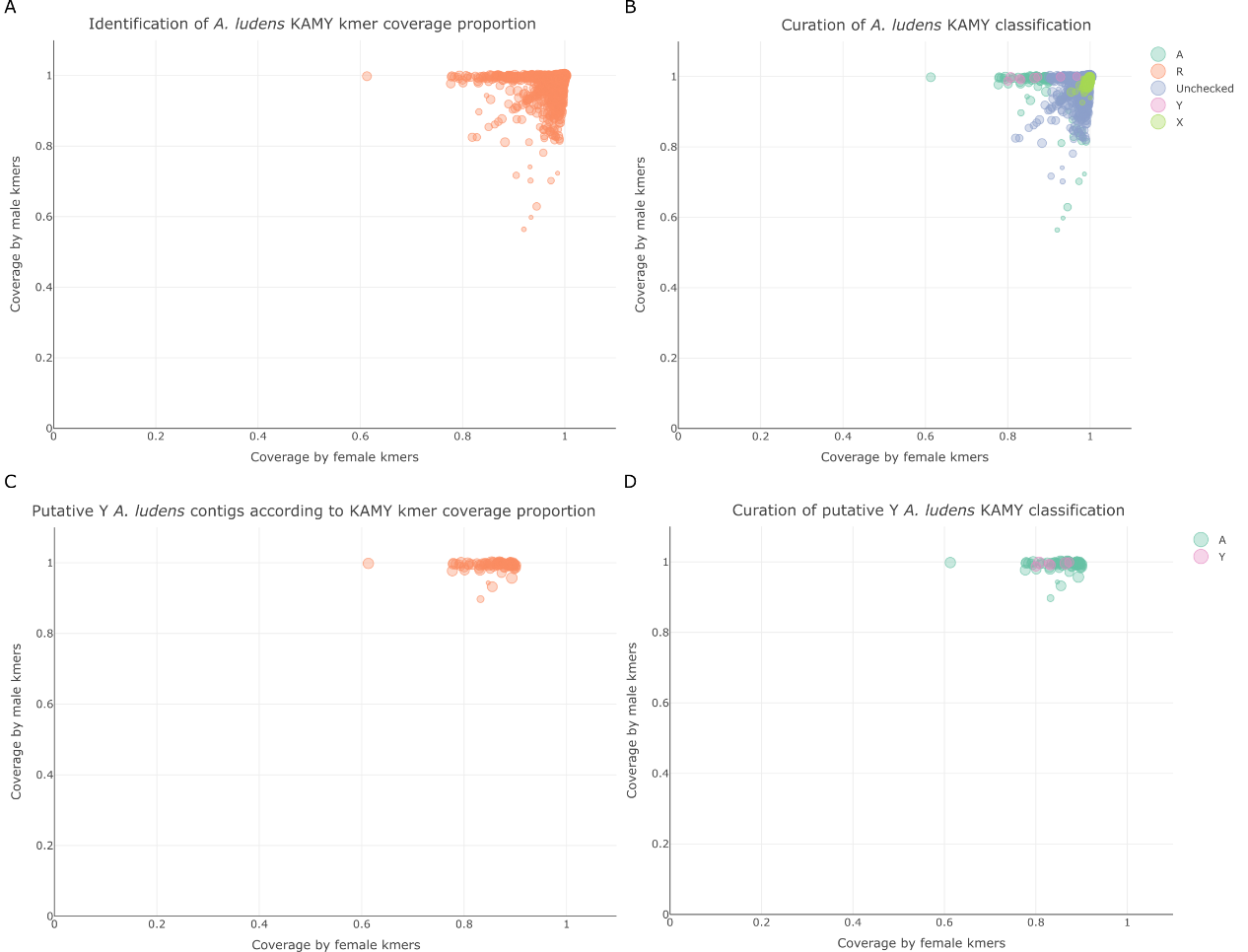

Figure S. 8 Scatterplot of male VS female kmer coverage of A. ludens contigs as identified by KAMY. A) kmer coverage of total A. ludens contigs. B) Curation-based classification of total A. ludens contigs in relation to kmer coverage values. C) kmer coverage of putative Y A. ludens contigs. D) Curation of putative Y contigs identified by KAMY.

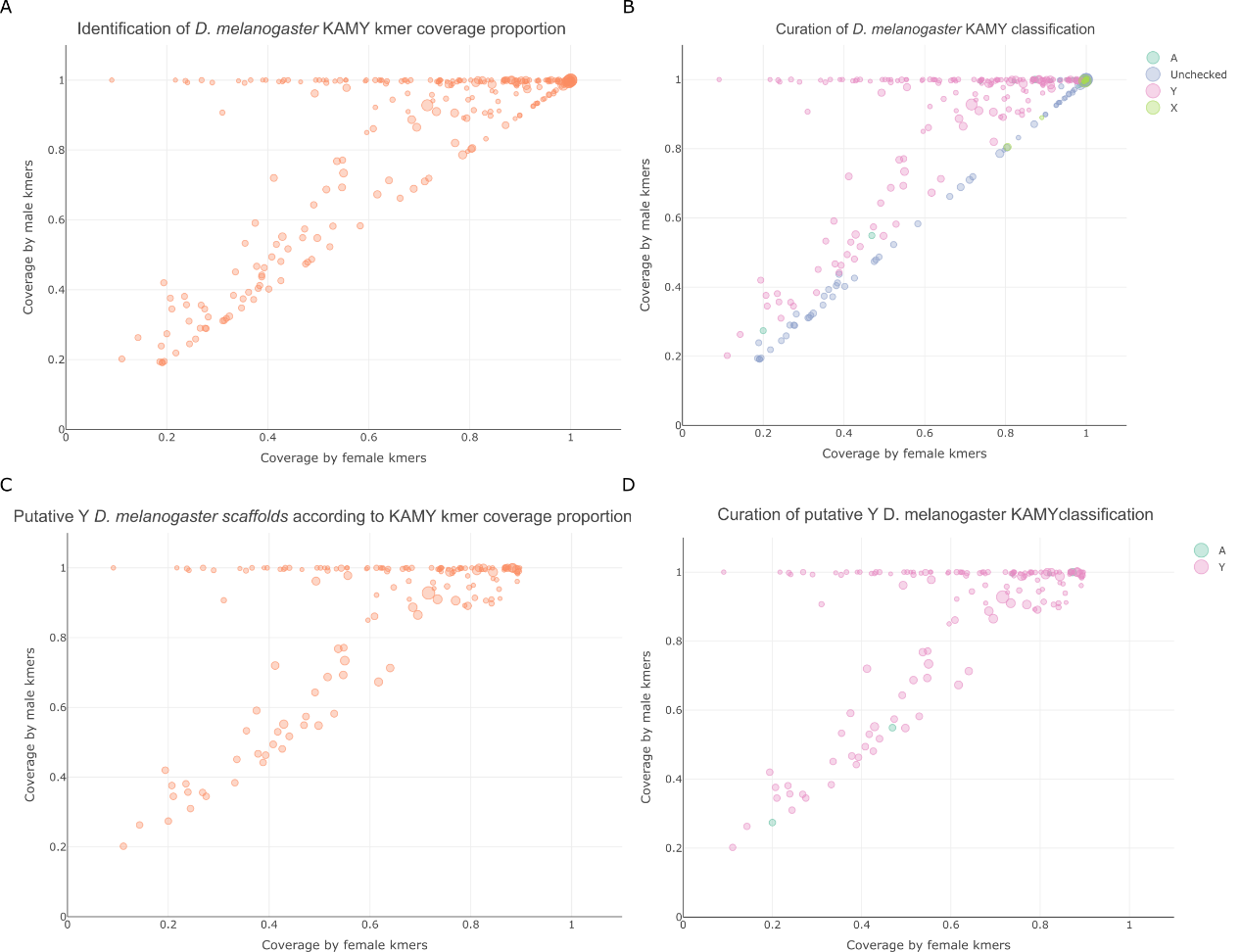

Figure S. 9 Scatterplot of male VS female kmer coverage of D. melanogaster scaffolds as identified by KAMY. A) kmer coverage of total D. melanogaster scaffolds. B) Curation-based classification of total D. melanogaster scaffolds in relation to kmer coverage values. C) kmer coverage of putative Y D. melanogaster scaffolds. D) Curation of putative Y contigs identified by KAMY.

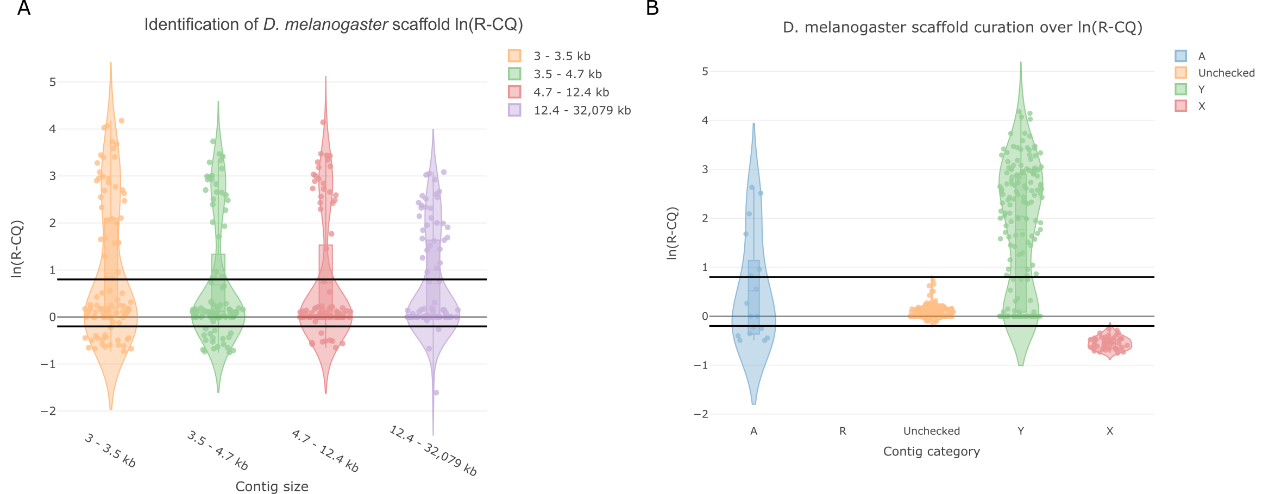

Figure S. 10 Violin plots of ln(R-CQ) values for D. melanogaster scaffolds, horizontal line indicates the ln=0.8 and ln= -0.2 cutoff used for indicating Y and X scaffolds respectively. A) Scaffolds are separated into four size-based quartiles. B) Distribution of ln(R-CQ) values over different curation categories.

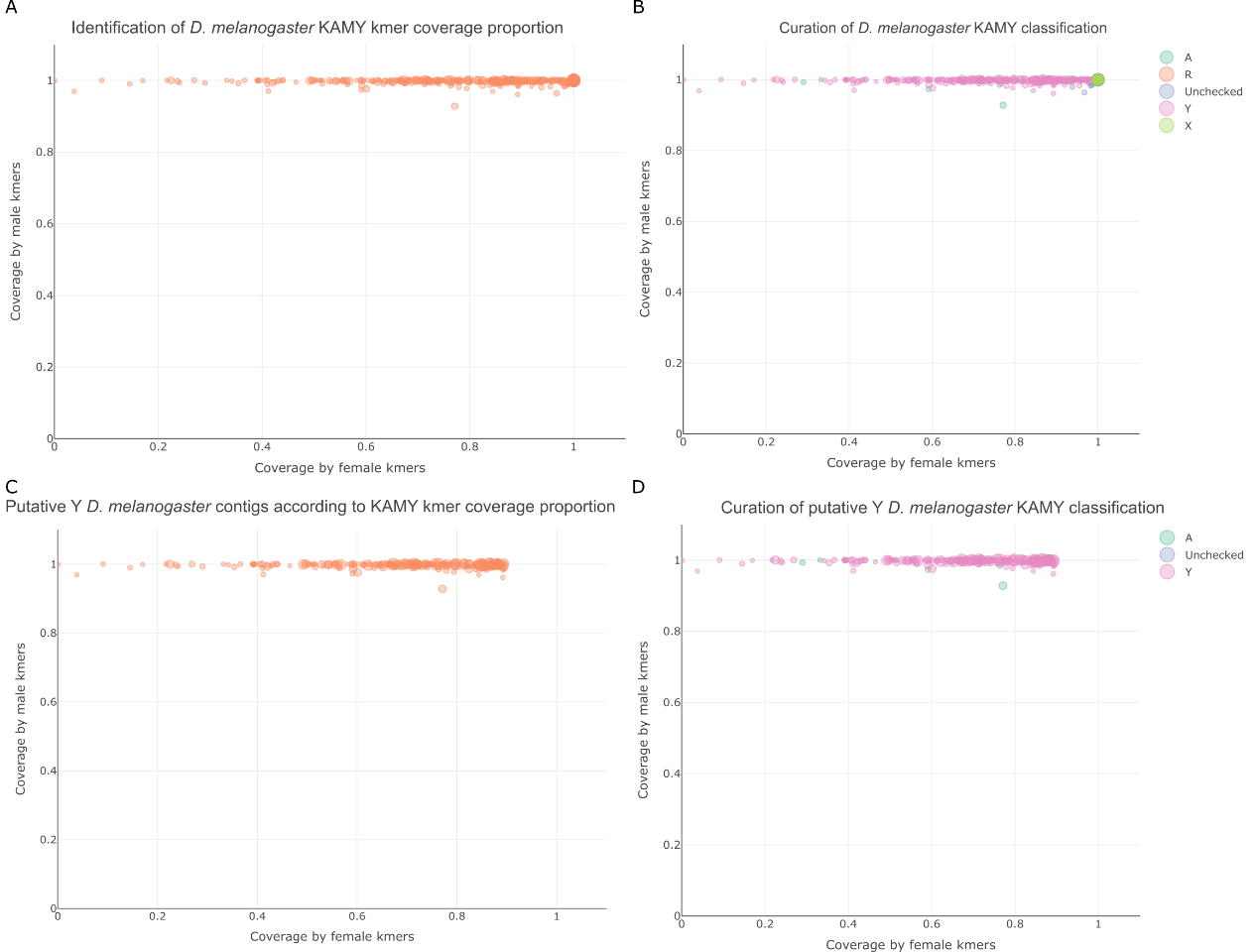

Figure S. 11 Scatterplot of male VS female kmer coverage of D. melanogaster contigs as identified by KAMY. A) kmer coverage of total D. melanogaster contigs. B) Curation-based classification of total D. melanogaster contigs in relation to kmer coverage values. C) kmer coverage of putative Y D. melanogaster contigs. D) Curation of putative Y contigs identified by KAMY.

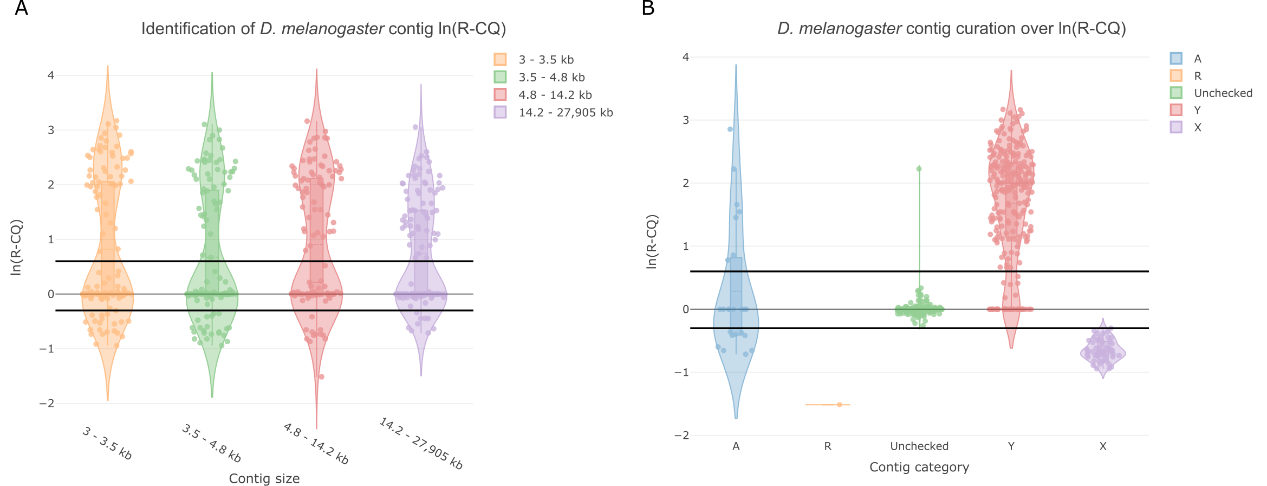

Figure S. 12 Violin plots of ln(R-CQ) values for D. melanogaster contigs, horizontal line indicates the ln=0.6 and ln= -0.3 cutoff used for indicating Y and X contigs respectively. A) Contigs are separated into four size-based quartiles. B) Distribution of ln(R-CQ) values over different curation categories.

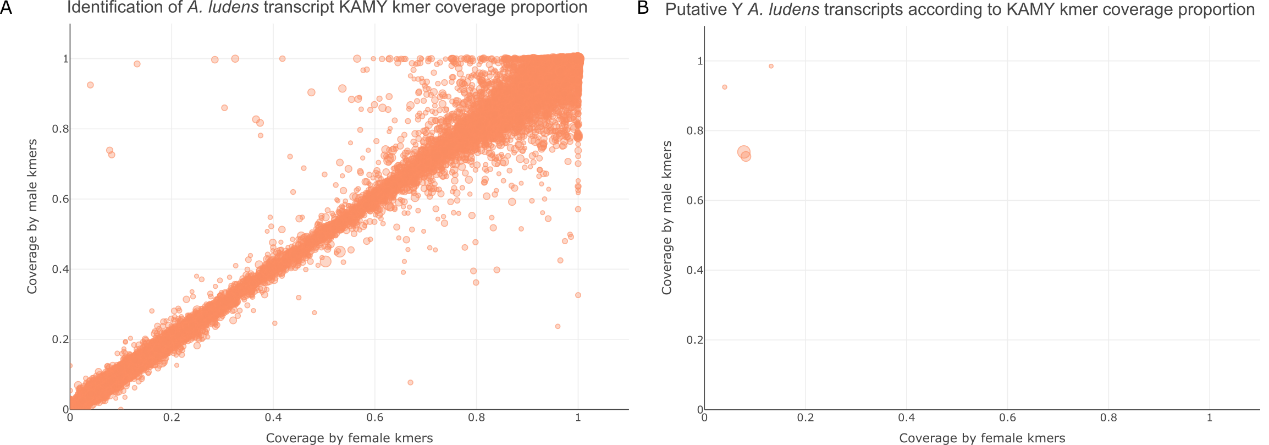

Figure S. 13 Scatterplot of male VS female kmer coverage of A. ludens testes transcripts as identified by KAMY. A) kmer coverage of total A. ludens testes transcripts. B) kmer coverage of putative Y A. ludens testes transcripts

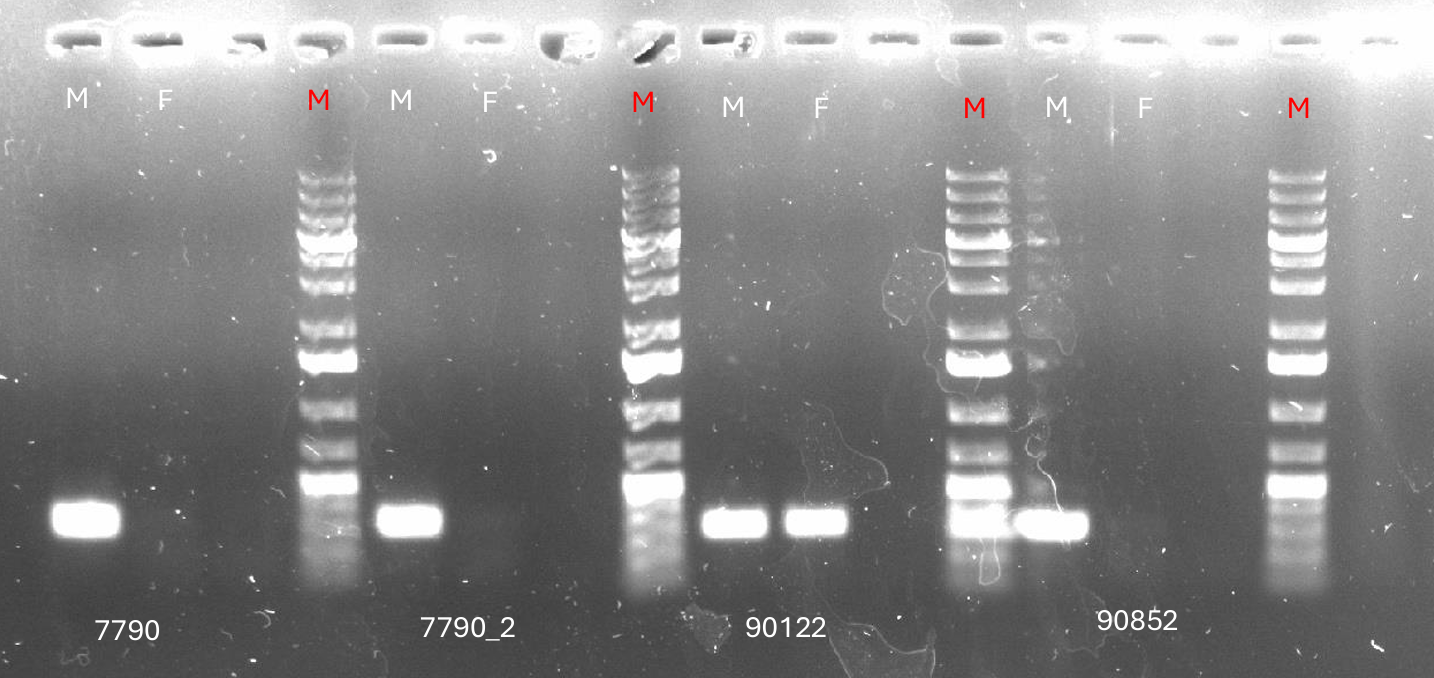

Figure S. 14 PCR validation of the putative Y transcripts of A. ludens 7790, 7790_2, 90122 and 90852. Line M and F contain the male and female specific amplification product while red M contains the molecular weight Marker.

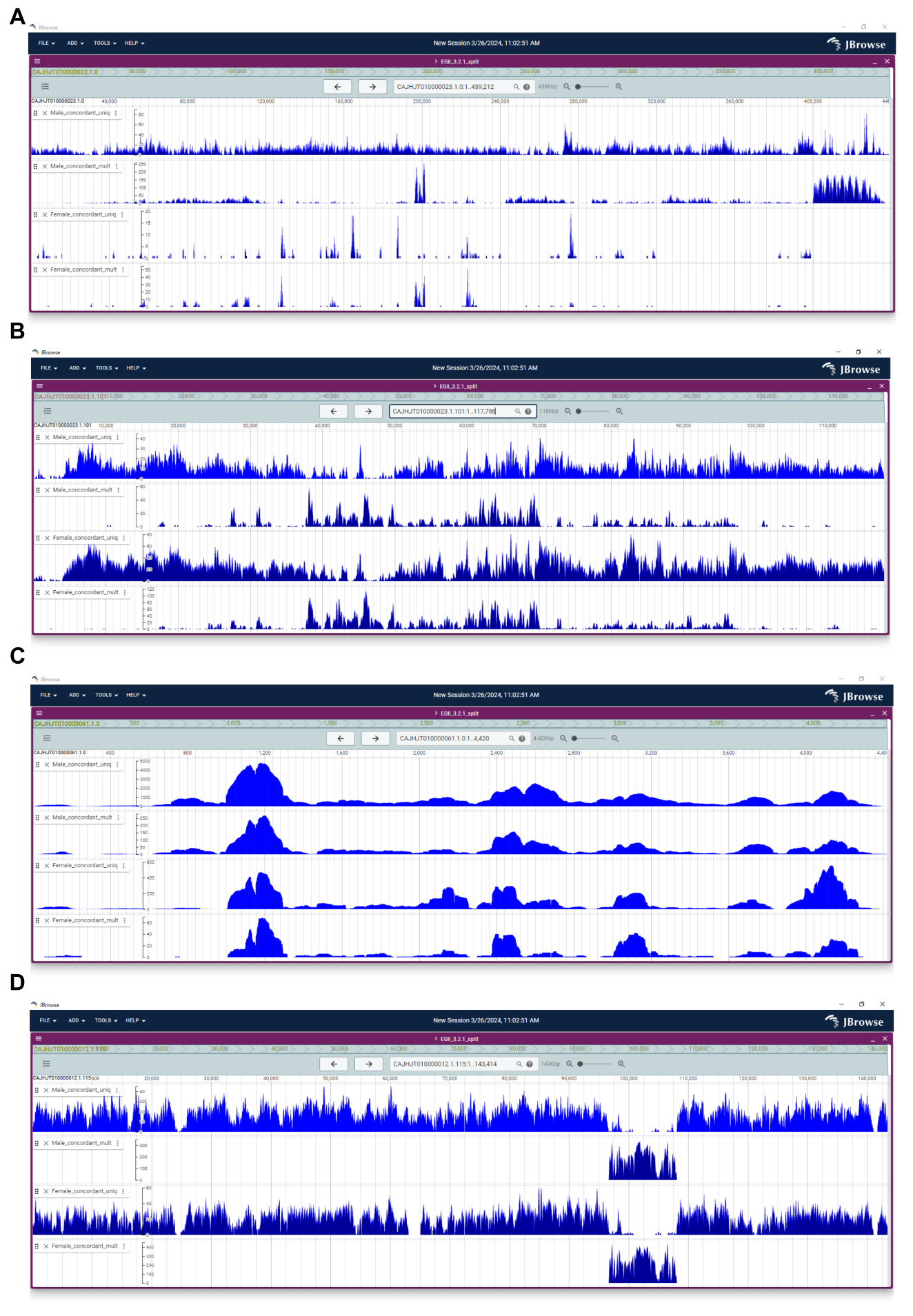

Figure S. 15 Screen snips of the male and female coverage plots from the Genome Browser of C. capitata. A) Example of a Y contig. B) Example of a false positive Autosomal contig. C) Example of a contig of the R (repeats) category. D) Example of an X contig.

### Supplementary tables

Table 1 Curation of R-CQ and KAMY output over C. capitata split genome.

| Curation | *R-CQ* | *Unique  R-CQ* | *R-CQ size (bp)* | *KAMY* | *Unique  KAMY* | *KAMY size (bp)* | *Total contigs* | | *Total size (bp)* |
| --- | --- | --- | --- | --- | --- | --- | --- | --- | --- |
| *Y* | *103* | *8* | *24,534,967* | *104* | *8* | *25,301,386* | *111* | *26,938,097* | |
| *A* | *0* | *0* | *0* | *29* | *29* | *6,255,452* | *29* | *6,255,452* | |
| *R* | *1* | *1* | *4,420* | *0* | *0* | *0* | *1* | *4,420* | |

Table 2 Curation of R-CQ and KAMY output over B. dorsalis split genome.

| Curation | *R-CQ* | *Unique  R-CQ* | *R-CQ size (bp)* | *KAMY* | *Unique  KAMY* | *KAMY size (bp)* | *Total contigs* | | *Total size (bp)* |
| --- | --- | --- | --- | --- | --- | --- | --- | --- | --- |
| *Y* | *2* | *0* | *4,265,511* | *2* | *0* | *4,265,511* | *2* | *4,265,511* | |
| *A* | *19* | *17* | *2,001,687* | *138* | *136* | *54,144,519* | *155* | *56,132,552* | |
| *R* | *0* | *0* | *0* | *0* | *0* | *0* | *0* | *0* | |

Table 3 Curation of R-CQ and KAMY output over B. zonata split genome.

| Curation | *R-CQ* | *Unique  R-CQ* | *R-CQ size (bp)* | *KAMY* | *Unique  KAMY* | *KAMY size (bp)* | *Total contigs* | | *Total size (bp)* |
| --- | --- | --- | --- | --- | --- | --- | --- | --- | --- |
| *Y* | *10* | *4* | *7,665,638* | *9* | *3* | *7,291,293* | *13* | *7,938,621* | |
| *A* | *14* | *13* | *1,148,655* | *6* | *5* | *1,432,708* | *19* | *2,572,363* | |
| *R* | *5* | *5* | *306,181* | *0* | *0* | *0* | *5* | *306,181* | |

Table 4 Curation of R-CQ and KAMY output over A. ludens split genome.

| Curation | *R-CQ* | *Unique  R-CQ* | *R-CQ size (bp)* | *KAMY* | *Unique  KAMY* | *KAMY size (bp)* | *Total contigs* | | *Total size (bp)* |
| --- | --- | --- | --- | --- | --- | --- | --- | --- | --- |
| *Y* | *4* | *2* | *442,059* | *2* | *0* | *344,782* | *4* | *442,059* | |
| *A* | *11* | *3* | *387,770* | *78* | *70* | *5,961,938* | *81* | *6,001,642* | |
| *R* | *0* | *0* | *0* | *0* | *0* | *0* | *0* | *0* | |

Table 5 Curation of R-CQ and KAMY output over D. melanogaster split genome.

| Curation | *R-CQ* | *Unique  R-CQ* | *R-CQ size (bp)* | *KAMY* | *Unique  KAMY* | *KAMY size (bp)* | *Total contigs* | | *Total size (bp)* |
| --- | --- | --- | --- | --- | --- | --- | --- | --- | --- |
| *Y* | *209* | *25* | *4386557* | *205* | *47* | *4237210* | *250* | *4733983* | |
| *A* | *7* | *5* | *51058* | *11* | *4* | *77986* | *15* | *91545* | |
| *R* | *0* | *0* | *0* | *0* | *0* | *0* | *0* | *0* | |

Table 6 Curation of R-CQ and KAMY output over D. melanogaster scaffolded genome.

| Curation | *R-CQ* | *Unique  R-CQ* | *R-CQ size (bp)* | *KAMY* | *Unique  KAMY* | *KAMY size (bp)* | *Total contigs* | | *Total size (bp)* |
| --- | --- | --- | --- | --- | --- | --- | --- | --- | --- |
| *Y* | *111* | *45* | *5149985* | *133* | *41* | *5301936* | *162* | *5803007* | |
| *A* | *6* | *4* | *38698* | *5* | *8* | *55674* | *10* | *72451* | |
| *R* | *0* | *0* | *0* | *0* | *0* | *0* | *0* | *0* | |

Table 7 Curations and total size of R-CQ X chromosome outputs against the genomes of assessed species. Abbreviations stand for Ceratitis capitata (Cc), Bactrocera dorsalis (Bd), Bactrocera zonata (Bz), Anastrepha ludens (Al) and Drosophila melanogaster (Dm).

| Curation | *X* | *A* | *R* |
| --- | --- | --- | --- |
| *Cc* | *125* | *1* | *0* |
| *Cc size* | *51,627,672 bp* | *256,468 bp* | *0 bp* |
| *Bd* | *7* | *15* | *4* |
| *Bd size* | *42,330,237 bp* | *1,099,363 bp* | *151209 bp* |
| *Bz* | *28* | *25* | *4* |
| *Bz size* | *39,337,604 bp* | *2,538,297 bp* | *529,213 bp* |
| *Al* | *342* | *200* | *2* |
| *Al size* | *25,598,582 bp* | *9,806,944 bp* | *100,975 bp* |
| *Dm (split)* | *54* | *9* | *1* |
| *Dm (split) size* | *23,608,662 bp* | *42,613 bp* | *14,179 bp* |
| *Dm (scaffolded)* | *42* | *7* | *0* |
| *Dm (scaffolded) size* | *23,740,909 bp* | *24,439 bp* | *0 bp* |

Table 8 Primers used for PCR validation of Y-linkage from C. capitata contigs.

| *Primer name* | *Sequence* |  | *Validated region* |
| --- | --- | --- | --- |
| *CcYG1_P1_F* | ACACATAATCCTTATTCTCTTGTATACGA |  | CAJHJT010000023.1.0 |
| *CcYG1_P1_R* | TATTCCCGCTCCCTTGTTTGTTT |  |  |
| *CcYG3_P3_F* | GGTCATTAACTGTTTGCATCCCC |  | CAJHJT010000023.1.6 |
| *CcYG3_P3_R* | AACCGCCTTAAAAATTCGTCTGG |  |  |
| *CcYG5_P1_F* | TCTGCATTTGCTCATTTATGGGG |  | CAJHJT010000023.1.73 |
| *CcYG5_P1_R* | TCCACAAGAACCGCCTTACTAC |  |  |
| *CcYG8_P1_F* | CTTACCGTCGTATTTGGGTAGTTTATT |  | CAJHJT010000023.1.73 |
| *CcYG8_P1_R* | CCGCTGTGCCGCTAATTTCG |  |  |
| *CcYG12_P2_F* | AAGAAAGACACAATTGATTTTTCTGA |  | CAJHJT010000023.1.152 |
| *CcYG12_P2_R* | TCTCTTCAACACTCTATCCAGCT |  |  |
| *CcYG13_P2_F* | TTCCACGATTTCCAGCATGTCTA |  | CAJHJT010000023.1.153 |
| *CcYG13_P2_R* | ACCAGAAGACAAAAGCAACACAA |  |  |
| *CcYG14_P2_F* | CTTCTTGTTTCTGCAACTACGGC |  | CAJHJT010000023.1.168 |
| *CcYG14_P2_R* | TTCCATGGTATTTAGGGAGAGAGT |  |  |
| *CcYG17_P1_F* | TCCTTTCAAAAAATTAATGCACAAA |  | CAJHJT010000023.1.168 |
| *CcYG17_P1_R* | CGATATTGTTCTTCCAATTTGATTCTA |  |  |
| *CcYG18_P2_F* | CCGACATTAACACGTGAGCATTT |  | CAJHJT010000023.1.168 |
| *CcYG18_P2_R* | ACCTCCACGTACAAATGCCATAA |  |  |
| *CcYG19_P1_F* | GCGTTAACTTGTATATACAAAGG |  | CAJHJT010000023.1.168 |
| *CcYG19_P1_R* | TCCAATTTGATTCTAAACGCTTT |  |  |
| *CcYG20_P2_F* | GCATTAACACGTGAGCATTTGGA |  | CAJHJT010000023.1.168 |
| *CcYG20_P2_R* | GAGAACCACTTCATTTGCCAAGG |  |  |
| *CcYG22_P1_F* | CCCCAACCATCTCTTCATTTTGG |  | CAJHJT010000023.1.168 |
| *CcYG22_P1_R* | TTGCGAGATAGTGATATGTGTGG |  |  |
| *CcYG23_P2_F* | CTTTCAAAGGGGGAAATTTCTGCT |  | CAJHJT010000023.1.200 |
| *CcYG23_P2_R* | TGAGTCAACACCAGCCACTTAAT |  |  |
| *CcYG24_P2_F* | CCATCGCAGGAGGGAAATAGACA |  | CAJHJT010000023.1.201 |
| *CcYG24_P2_R* | ACGATTTTCGACGCCATAATCCT |  |  |

Table 9 Primers used for PCR validation of Y-linkage from B. dorsalis contigs.

| *Primer name* | *Sequence* |  | *Validated region* |
| --- | --- | --- | --- |
| *Bd_3025_F* | TTCGCCTGGTTTAGATTCG |  | Scaffold_6.8 |
| *Bd_302_R* | TCTGTGAACTGCTGGATTG |  |  |
| *Bd_0888_F* | AGTTATGCTGCCAGATCGTAGG |  | Scaffold_6.8 |
| *Bd_0888_R* | GAAATCACGCTTAACTCTCGGC |  |  |
| *Bd_5507_F* | CCGCTATCACAAGACAAGAC |  | Scaffold_6.8 |
| *Bd_5507_R* | TTTGGAGTAAGGAACCTGTTG |  |  |
| *Bd_3132_F* | GGATGAAGCACCAAGAAGAG |  | Scaffold_6.9 |
| *Bd_3132_R* | CTCACTCAGCTCACCATAAAG |  |  |
| *Bd_3522_F* | CGTCATGATCTGGAGTCCGG |  | Scaffold_6.8 |
| *Bd_3522_R* | GACACTTGGGGGCGTATACC |  |  |

#### Supplementary methods

#### PCR validation

A set of curated Y contigs from *C.* *capitata* were selected for PCR validation. Primer design sought to generate PCR products of 400 to 1,000 bp and was assessed for its uniqueness using the Primer BLAST. Template DNA was derived from males and females individually using pools of 4 insects so that multiple haplotypes can be captured. Virgin *C. capitata* insects were collected from the “EGII laboratory” populations reared at the University of Thessaly. Genomic DNA was extracted using the protocol described in Calderón-Cortés et al. 2010 and quantified using a Quawell Q3000 spectrophotometer so that 30 ng/µl were used in each reaction. PCR reactions were conducted using a KAPA Taq (Merck) according to manufacturer’s protocol. PCR reactions were carried out at a Bio-Rad T100 thermal cycler and the PCR products were visualized on 1 % agarose gel. When a band was present in only males it deemed that the primer pair was from a Y-linked contig.
